# Third generation sequencing revises the molecular karyotype for *Toxoplasma gondii* and identifies emerging copy number variants in sexual recombinants

**DOI:** 10.1101/2020.03.10.985549

**Authors:** Jing Xia, Aarthi Venkat, Michael L. Reese, Karine Le Roch, Ferhat Ay, Jon P. Boyle

## Abstract

*Toxoplasma gondii* is an obligate intracellular parasite that has a significant impact on human health, especially in the immunocompromised. This parasite is also a useful genetic model for intracellular parasitism given its ease of culture in the laboratory and relevant animal models. However, as for many other eukaryotes, the *T. gondii* genome is incomplete, containing hundreds of sequence gaps due to the presence of repetitive and/or uncloneable sequences that prevent complete telomere-to-telomere de novo chromosome assembly. Here, we report the first use of single molecule DNA sequencing to generate near complete de novo genome assemblies for *T. gondii* and its near relative, *N. caninum*. Using the Oxford Nanopore Minion platform, we dramatically improved the contiguity of the *T. gondii* genome (N50 of ∼6.6Mb) and increased overall assembled sequence compared to current reference sequences by ∼2 Mb. Multiple complete chromosomes were fully assembled as evidenced by clear telomeric repeats on the end of each contig. Interestingly, for all of the *Toxoplasma gondii* strains that we sequenced (RH, CTG, II×III F1 progeny clones CL13, S27, S21, and S26), the largest contig ranged in size between 11.9 and 12.1 Mb in size, which is larger than any previously reported *T. gondii* chromosome. This was due to a repeatable and consistent fusion of chromosomes VIIb and VIII. These data were further validated by mapping existing *T. gondii* ME49 Hi-C data to our assembly, providing parallel lines of evidence that the *T. gondii* karyotype consists of 13, rather than 14, chromosomes. In addition revising the molecular karyotype we were also able to resolve hundreds of repeats derived from both coding and non-coding tandem sequence expansions. For well-known host-targeting effector loci like rhoptry protein 5 (ROP5) and ROP38, we were also able to accurately determine the precise gene count, order and orientation using established assembly approaches and the most likely primary sequence of each using our own assembly correction scripts tailored to correcting homopolymeric run errors in tandem sequence arrays. Finally, when we compared the *T. gondii* and *N. caninum* assemblies we found that while the 13 chromosome karyotype was conserved, we determined that previously unidentified large scale translocation events occurred in *T. gondii* and *N. caninum* since their most recent common ancestry.

## INTRODUCTION

*Toxoplasma gondii* and its Apicomplexan relatives are highly successful animal pathogens, infecting a wide variety of warm-blooded animals including humans and domesticated animals. *T. gondii* infection can lead to severe toxoplasmosis in immunocompromised individuals and in congenitally-infected fetuses (Joynson and Wreghitt 2005), and is a leading cause of blindness due to its ability to infect the eye causing ocular toxoplasmosis (Jones and Holland 2010). *T. gondii* belongs to the phylum Apicomplexa, a large group of animal and human pathogens including *Neospora*, *Eimeria*, *Plasmodium* and *Cryptosporidium*. The ease of genetic manipulation, accessibility to cellular and biochemical experiments, and well-established animal model make *T. gondii* an important system for studying the biology of Apicomplexans (Kim and Weiss 2004). Genomic analysis tools for this organism have been under development for decades. Data housed at ToxoDB.org, the primary genomic repository for *T. gondii* genome-wide data, presently includes sequence, *de novo* assemblies and annotation of multiple *T. gondii* genomes, next-generation sequence data for an additional 60 *T. gondii* genomes, as well as draft assemblies for both *Hammondia hammondi* and *Neospora caninum* (Lorenzi *et al*. 2016a).

Availability of a complete reference genome that contains accurate representations of all small- or large-scale structural variants is essential to have a better understanding of gene content, genotype-phenotype relationships, and the evolution of unique traits in parasites of humans and other animals. However, like all eukaryotic genomes, a substantial part of the *T. gondii* genome consists of repetitive elements (Matrajt *et al*. 1999), making gap-free *de novo* assembly impossible using standard 1st or 2nd generation sequencing approaches. Even with exceptionally high coverage, these approaches fail to resolve repetitive regions or complex structural variants with repeat units that are larger than the size of the individual reads. Three of the *T. gondii* reference genomes in ToxoDB (Gajria *et al*. 2008) were constructed by combining high quality 1st generation Sanger (Sanger *et al*. 1977) and 2nd generation 454 (Roche Applied Science) sequence data, yet these genomes still have hundreds of sequence gaps of unknown sequence content and length. Assembly gaps typically mask repetitive regions which can contain previously unknown protein-coding genes, additional copies of genes found in tandem gene arrays, and in some cases, they may also lead to incorrect predictions of chromosomal structure. This problem is not unique to *T. gondii* and other apicomplexan genomes. For example, all versions of the human genome have thousands of gaps due to incorrect assembly of repetitive sequence (Vollger *et al*. 2019), including assemblies generated recently using new sequencing technologies like those applied here.

In recent years single molecule sequencing approaches (developed by Oxford Nanopore and PacBio) have revolutionized *de novo* sequence assembly by enabling high-throughput generation of kilobase-sized sequence reads. These approaches have allowed for resolution of the vast majority of repeat-driven sequence assembly gaps, the detection and assembly of previously intractable structural variants with species, and when combined with 2nd generation sequencing data can be used to generate near-complete *de novo* genome assemblies with high (>99%) nucleotide accuracy. Indeed, whole genome assemblies of several organisms including bacteria (Madoui *et al*. 2015; Fournier *et al*. 2017; Diaz-Viraque *et al*. 2018), parasites (Lapp *et al*. 2018), plants (Schmidt *et al*. 2017; Michael *et al*. 2018), and mammals (Jain *et al*. 2018) using a such a hybrid approach have been reported, generating assemblies of unprecedented contiguity. Here, we apply Oxford Nanopore sequencing and de novo assembly using the Minion Platform to multiple isolates of *T. gondii*, F1 progeny of a cross between two canonical *T. gondii* strains, and one of its nearest extant relatives, *Neospora caninum*. Using these data, we have generated mostly gap-free, telomere-to-telomere assemblies for the majority of the *T. gondii* chromosomes from each sequenced isolate and/or species. These assemblies revise the molecular karyotype for *T. gondii*, formally establishing that this parasite has 13, rather than 14, chromosomes. This result was confirmed by Hi-C sequencing technologies as described previously (Bunnik *et al*. 2019).This karyotype is conserved across all *T. gondii* parasite strains that we sequenced, and is also conserved in *N. caninum*. Our new assemblies have also resolved copy number estimates at simple- and complex repetitive loci, including multiple tandem gene arrays harboring known host-targeting effector genes. Moreover, by directly assembling genomes from multiple F1 progeny experimental crosses, we have determined for the first time that changes in gene copy number occur during sexual recombination, and that this occurs frequently at some *T. gondii* loci (like the ROP5 locus) and infrequently at others (like the MAF1 locus), and may explain some of the differences in pathogenicity between *T. gondii* strains. Finally, our *de novo* assembly of the *N. caninum* Liverpool genome shows that the current draft assembly of this organism dramatically overestimates chromosome-level synteny with *T. gondii*. This discovery is also corroborated in multiple *N. caninum* strains in a companion paper (See cover letter for details on co-submitted manuscript).

## RESULTS

### *De novo* assembly of *Tg*RH88 genome using nanopore reads revises the *T. gondii* karyotype

The majority of the *T. gondii* isolates collected from North America and Europe belong to three predominant clonal lineages, types I, II, and III (Sibley and Ajioka 2008) and RH strain is a representative strain of the type I lineage (Pfefferkorn and Pfefferkorn 1976). While RH strain shows some unique phenotypes including higher growth rate *in vitro* and inability to form cysts (Villard *et al*. 1997; Khan *et al*. 2009) and is most frequently used in *T. gondii* studies, a complete *de novo* assembly for *Tg*RH genome is lacking. Therefore, our initial efforts were to use long read single molecule sequencing to generate a more complete *Tg*RH88 assembly.

To take advantage of the long-read technology, high-molecular weight genomic DNA of *Tg*RH88 strain was extracted using an optimized protocol which was originally designed for extraction of gram-negative bacteria and mammalian cell DNA (Quick 2018). This protocol included a phenol-chloroform-isoamyl alcohol extraction followed by ethanol precipitation, and the use of large bore pipette tips for all manipulations (see Materials and Methods). For TgRH88, we obtained 24 μg of genomic DNA from 2×10^8^ tachyzoites, and 400 ng was used for MinION sequencing. A 48-hour sequencing run on a single flow cell yielded 648,491 reads containing 7.40 Gb of sequences for *Tg*RH88 genome. Assuming a 65 Mb *T. gondii* genome size, these reads represented a genome coverage of ∼114×. While more detailed sequence metrics are described below, the sequence reads we obtained robustly aligned to the TgGT1 reference genome found at ToxoDB.org (**Figure 1A**).

**Figure 1.**
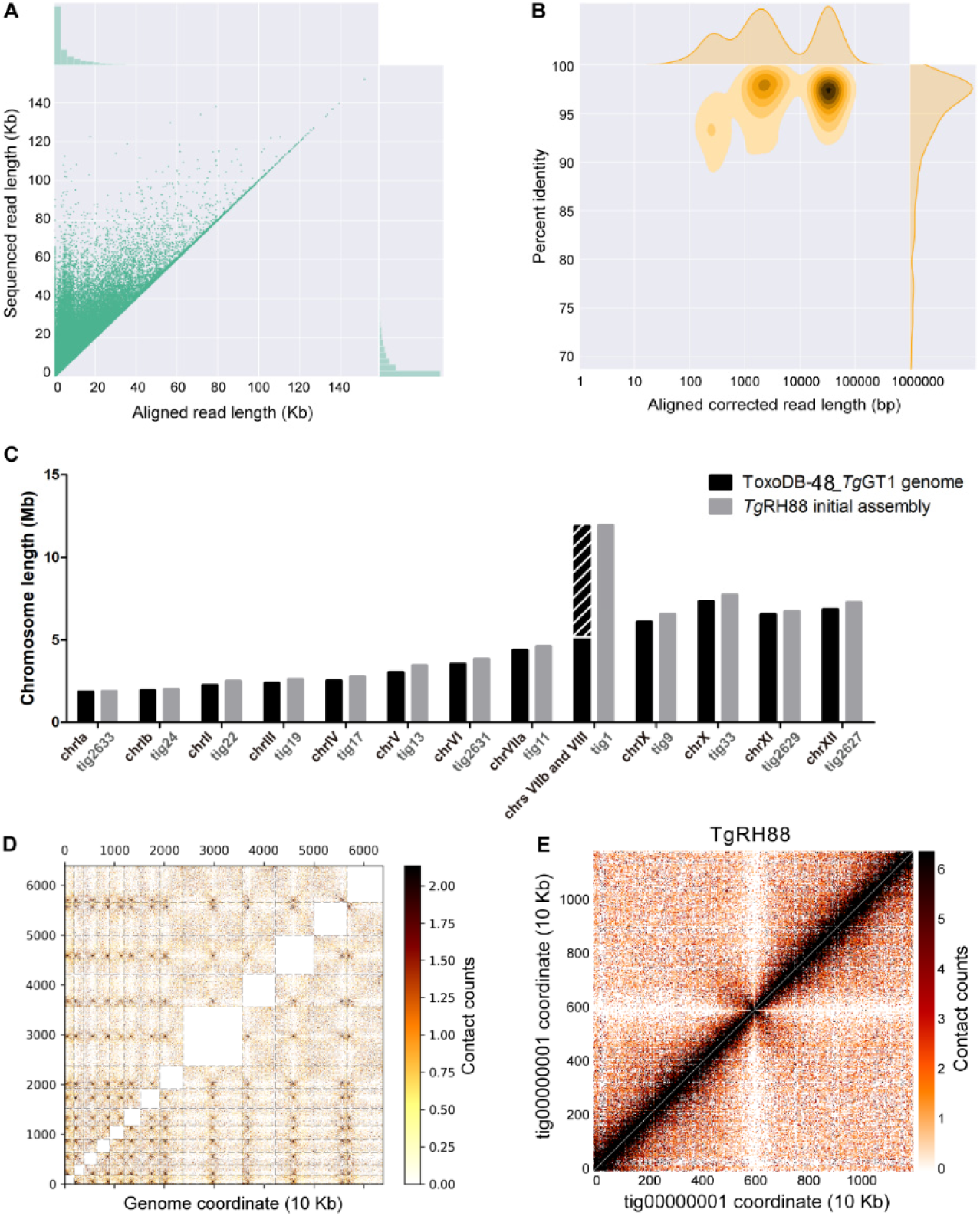
Primary de novo assembly of TgRH88 genome using nanopore reads revises T. gondii karyotype. (A) Bivariate plot showing a comparison of the aligned read length with the sequencing read length. (B) Bivariate plot showing a comparison of the aligned corrected read length (log10 transformed) with the percent identity. In this case corrected reads refer to the method deployed by Canu using read overlap. (C) Histogram showing comparison of chromosome size between ToxoDB-48_TgGT1 genome and TgRH88 initial long-read assembly. (D) Interchromosomal Hi-C contact-count heat map plotted using the TgRH88 initial long-read assembly sequence showing 13 chromosomes in the assembly. (E) Intrachromosomal Hi-C contact-count heat map plotted using the sequence of TgRH88_tig00000001 in TgRH88 initial long-read assembly showing no aberrant signal along the contig

Before assembly, Canu v1.7.1 corrected all reads > 1000 bp, and after alignment we found that 99.97% of the corrected *Tg*RH88 reads could be mapped to the ToxoDB-48_*Tg*GT1 reference genome, and the mean percent identity was 92.9% (**Figure 1B**). The resulting Canu-corrected reads were subjected to *de novo* assembly using Canu v1.7.1, which yielded a *Tg*RH88 primary assembly with a size of 64.40 Mb. The assembly consisted of 23 contigs and 6 of them contained telomeric repeat sequences (TTTAGGG or CCCTAAA) on at least one end. Interestingly, by aligning the *Tg*RH88 assembly sequences to the ToxoDB-48_*Tg*GT1 reference genome, we noticed that the sequences annotated as chrVIIb and chrVIII were parts of a single contig in our *Tg*RH88 assembly (TgRH88_tig00000001; **Figure 1C**). This contig, which was 11.93 Mb in length, was longer than any previously reported *T. gondii* chromosome, and suggested to us that *T. gondii* consists of 13 (instead of 14) chromosomes, with the chrVIIb and chrVIII being actually a single chromosome. Given that prior work using Hi-C chromosome conformation capture sequencing suggested a similar fusion between chromosomes VIIb and VIII (Bunnik *et al*. 2019), we mapped the Hi-C reads from that study onto our TgRH88 de novo assembly to determine if it has similar contact counts. As shown in **Figure 1D** the Hi-C data identified the position of 13, rather than 14, interchromosomal contact points (representing centromeres; (Bunnik *et al*. 2019)) and an intrachromosomal contact map across TgRH88tig00000001 indicating that this did indeed represent a single contiguous chromosome (**Figure 1E**). These findings provide direct assembly-based evidence that sequence fragments previously referred to as distinct chromosomes (VIIb and VIII) were in fact two parts of the same chromosome. We have named this contig TgRH88_tig00000001_chrVIII.

The RH strain is one of the most commonly used laboratory strains given its genetic tractability and robust *in vitro* growth characteristics (Saeij *et al*. 2005), but at the genomic level this strain has been subject to much less formal annotation compared to strain types GT-1, ME49 and VEG. However, we did identify two additional RH strain de novo assemblies in Genbank, one generated using Illumina technology (Lau *et al*. 2016); Genbank BioProject PRJNA294483) and the other using single-molecule long read sequencing using PacBio RS technology (BioProject PRJNA279557). As expected, our Nanopore and the PacBio RS assembly were more contiguous than the de novo Illumina assembly, having similar contig N50 values (6.7 Mb for both; **Figure 2A**) and both predicting the ∼12 Mb chromosome VIII. Moreover total size of the predicted nuclear-encoded genomes in each assembly differ by only 0.8% (a raw difference of 513,977 bp of additional sequence present in the Nanopore assembly; discussed further below). Despite this congruence of the two long-read assemblies, the most striking observed difference between them is the total number of contigs (23 vs. 203, respectively) in each of the final assemblies (prior to scaffolding; **Figure 2A**). Based on Nucmer alignments, ∼362 Kb of this unplaced sequence is shared between the two assemblies, while the remaining sequences (∼70 and 2700 Kb for the Nanopore and PacBio RH assemblies, respectively) were unshared. We also used blast to identify contigs harboring what appeared to be sequence derived from either of the two organellar genomes of *T. gondii*, the apicoplast and the mitochondrion and compared them to the other existing RH assemblies (Figure 2A). One contig ∼35 Kb in size from each assembly was derived from the plastid genome, which is the expected size for this organellar genome (Lorenzi *et al*. 2016b). For the mitochondrial genome, we identified two non-chromosomal contigs of ∼15 Kb in size in our assembly that likely harbored at least fragments of this genome (Figure 2A), while there were no contigs in the other RH assemblies with any resemblance to a mitochondrial genome. It is likely that this is due to pre-filtering genome assemblies prior to submission to genbank to eliminate mitochondrial sequences. However for all of our Nanopore assemblies presented here and submitted to genbank, we have annotated all putative mitochondrial genome contigs with the “location=mitochondria” tag so that they would pass the filters in Genbank. However given the complexities of the mitochondrial genome and its assembly for *T. gondii* (as evidenced by recent work from the Kissinger group; https://www.biorxiv.org/content/10.1101/2020.05.16.099366v1), we did not perform any extensive analyses of these sequences.

**Figure 2:**
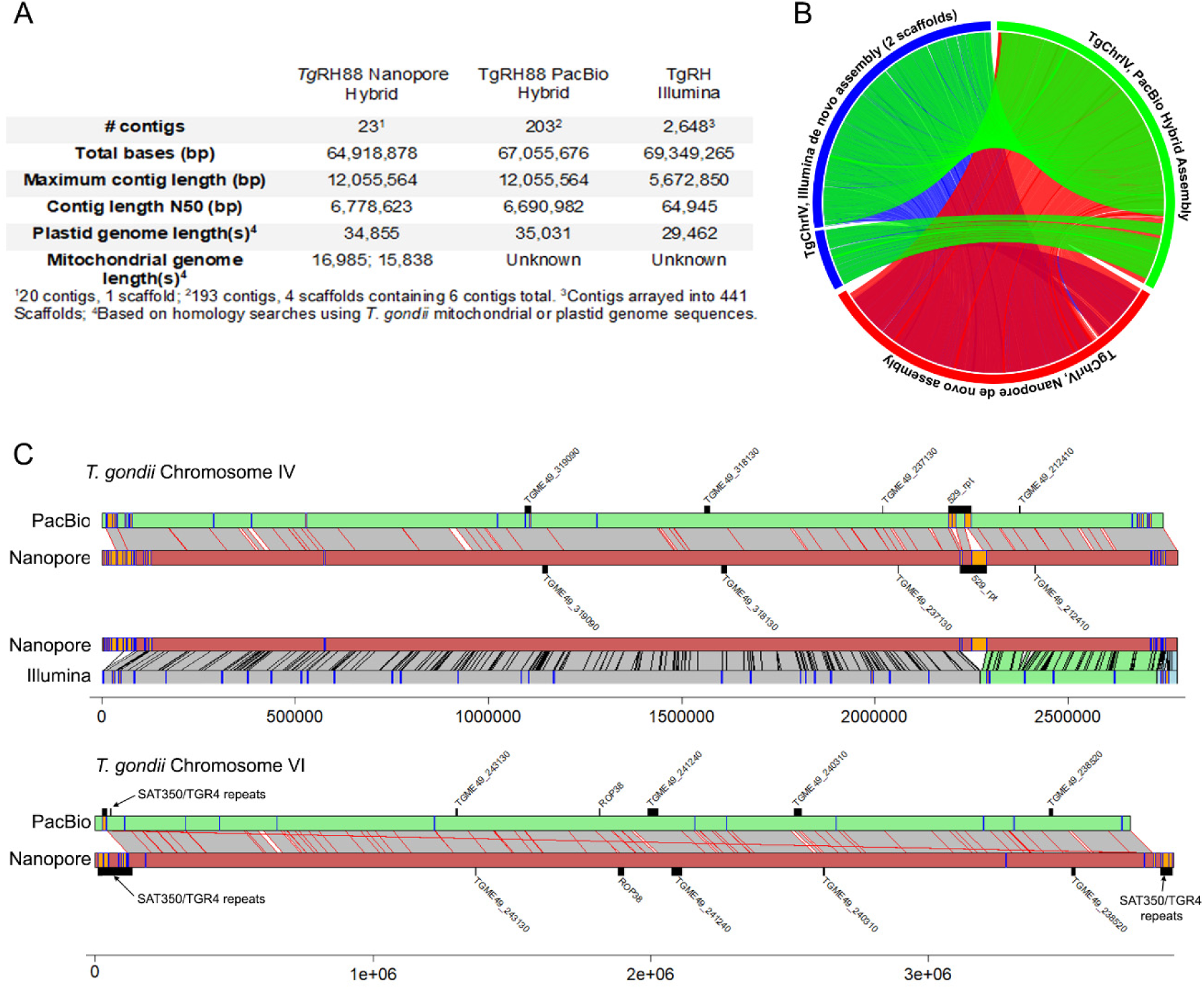
Comparisons between the current Nanopore assembly of T. gondii strain RH88 and an existing long read (using PacBio RS and Illumina technology) and short read (Illumina only) assemblies. A) Assembly statistics for each. B) Circos plot of Nucmer pairwise alignments across all 3 assemblies for T. gondii chromosome IV. All alignments > 10000 bp and > 90% identity are shown. C) Pairwise alignments for chromosomes IV (top) and VI (bottom) along with locations of select tandem repeats identified either de novo (orange or blue bars on the chromosome scaffolds) or known from prior studies (above or below chromosome scaffolds). For T. gondii chromosome IV comparisons to both the PacBio and Illumina assemblies are shown, while only the long-read comparison is shown for T. gondii chromosome VI.

To further compare our assembly to the two existing RH88 de novo assemblies we compared them using Mummer (Delcher *et al*. 2003) and Tandem Repeats Finder (Benson 1999). While complete chromosomes aligned with high synteny between the Nanopore and PacBio assemblies (Figure 2B), chromosomes in the Illumina only RH assembly were distributed across multiple scaffolds (e.g., see TgRH88 Chromosome IV; Figure 2B). Given that long read sequencing technologies improve sequence contiguity across tandem repeats compared to shorter read technologies like Illumina (especially at tandem repeats that are longer than the typical Illumina read), we mapped all tandem repeats with periods greater than 500 bp as well as select known repeats in the RH assemblies and mapped them along with nucmer-generated pairwise sequence alignments for chromosomes IV and VI (Figure 2C). For chromosome IV, the most striking difference between the long read assemblies and that generated using Illumina is the expanded assembly at the locus harboring the well-characterized 529 bp repeat (Figure 2C, top). We also found that loci harboring select tandem repeat genes (e.g., TGME49_318130 and TGME49_319090) were also expanded in the long-read *T. gondii* chromosome IV assemblies compared to that generated using solely Illumina sequences. Alignments for *T. gondii* chromosome VI are also shown in figure 2C, in this case between just the long read assemblies to illustrate differences and similarities. In multiple cases the subtelomeric repeat sequences SAT350 and TGR4 were found in larger arrays in the Nanopore assembly, as was the case for the known tandem rhoptry gene array encoding ROP38. In contrast gene TGME49_240310 was found in a larger array in the PacBio assembly. Overall these data indicate the utility of long read sequencing for resolving complex tandem repeats in *T. gondii*, especially those that are larger than typical 1^st^ or 2^nd^ generation sequence reads. Moreover with a few minor exceptions both existing long-read sequencing technologies give similar results for the size and complexity of these repeats, and results in remarkably similar estimates for nuclear chromosome and plastid genome sizes.

### Nanopore read quality assessment and *de novo* genome assembly statistics

To determine whether the chr VIIb/VIII fusion was unique to TgRH88 or was also present in other isolates, we sequenced and assembled genomes of *Tg*ME49, *Tg*CTG, *Tg*ME49×*Tg*CTG F1 progeny (CL13, S27, S21, S26, and D3X1), and *N. caninum* Liverpool strain (**Table 1**) by performing 6 MinION runs with 6 R9.4.1 flow cells. On average, a single flow cell yielded approximately 600,000 reads containing more than 7.8 Gb of sequences over a 48-hour run, and for each strain, total yield varied from 0.6 Gb to 7.4 Gb (**Table S2**). The read length N50s (the sequence length of the shortest read at half of the total bases) were longer than 18 Kb, and the maximum read lengths varied between 116 Kb and 266 Kb (**Table S2**). The average Phred quality scores (a measure of the quality of the base-call, representing the estimated probability of an error (Ewing and Green 1998; Ewing *et al*. 1998)) for all libraries were ∼10.0, except for the *Tg*RH88 library, whose average Phred score was 8.9 (**Table S2**). Aligning the reads against their most relevant “Reference” genome in ToxoDB (based on species and then closest genotype) revealed that 96% of the *T. gondii* reads and 85% of the *N. caninum* reads could be mapped, with a mean percent identity of ∼86% (**Table S2**). After reads were corrected using error-correction in Canu, 99% of the *T. gondii* reads and 98.07% of the *N. caninum* reads could be mapped to their reference genomes, and the mean percent identities were ∼95% (**Table S3**). This first round of error correction (based on alignment and overlap between Nanopore reads as implemented in Canu; (Koren *et al*. 2017)) was sufficient for making conclusions about previously unappreciated structural variation within and between species.

**Table 1.**
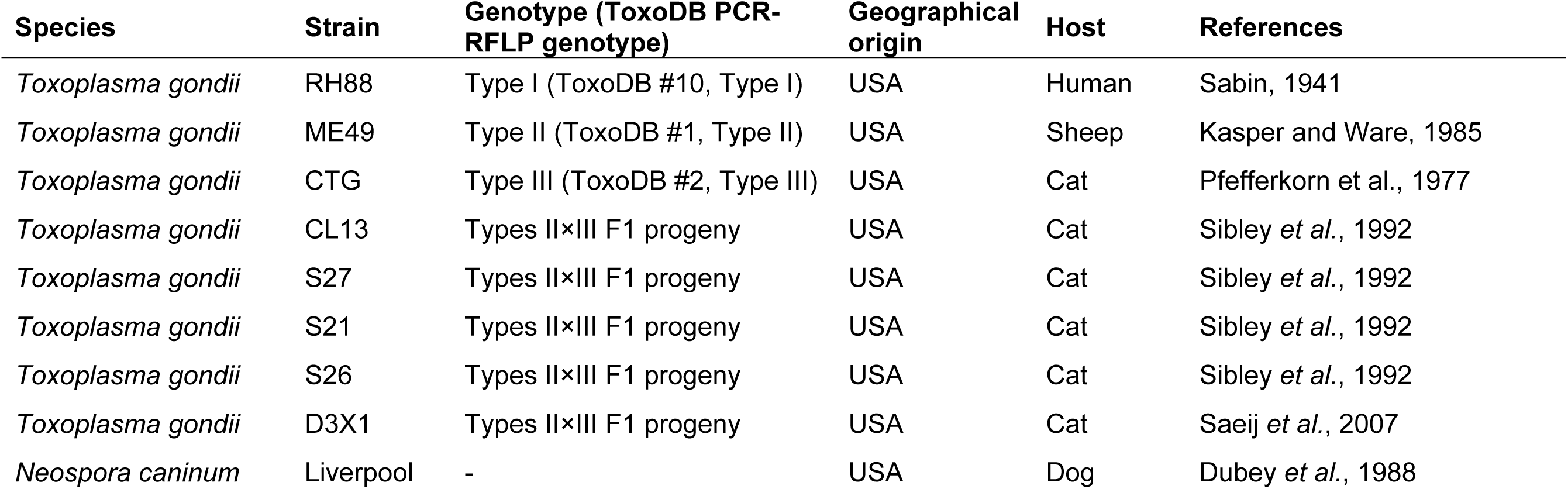
Description of the *T. gondii* and *N. caninum* strains sequenced in this study.

The primary assemblies had a median number of contigs of 38.5 for the *T. gondii* strains and 58 for the *N. caninum* Liverpool strain (**Table S4**). The median assembly size of the *T. gondii* strains was 64 Mb, with a median contig length N50 of 6.63 Mb, and a median L50 of 4 (**Table S4**). The *N. caninum* primary assembly had a size of 61.8 Mb, with a contig N50 size of 6.38 Mb and an L50 of 4 (**Table S4**). Depending on the strain, contigs containing plastid genome sequence ranged in size from 8 Kb to 214 Kb (Table 2), which is mostly inconsistent with the known size of the plastid genome. While for the 8Kb contig (from strain S26) was likely due to low sequence coverage, the larger contigs may be either concatenated versions of the plastid genome or hybrid assemblies of plastid sequence along with some other non-nuclear sequence. This suggests that the plastid genome may be more susceptible than the nuclear genome to artificial concatenations during assembly of error-prone long sequence reads, although we have not investigated this directly. In contrast to the organellar genomes the nuclear *T. gondii* genomes assembled with high consistency across multiple strain types and their F1 progeny. Although the mean percent identity of the aligned Canu-corrected reads to the reference genome (∼95%) was improved compared to the raw reads (∼86%), we used a second round of error correction to improve sequence accuracy. To do this, we mapped whole-genome Illumina paired-end reads (SRA: SRR5123638, SRR2068653, SRR5643140, or ERR701181) to the *Tg*RH88, *Tg*ME49, *Tg*CTG, or *Nc*Liv primary assemblies generated by Canu, and iteratively polished the assembly contigs four times using Pilon v1.23. The polished contigs were then reassembled using Flye v2.5. The resulting contigs/scaffolds were then assigned, ordered, and oriented to chromosomes using relevant ToxoDB-48 reference genomes (**Table 2 and Table S1**). These polished assemblies have been deposited in Genbank (under BioProject number PRJNA638608).

**Table 2.**
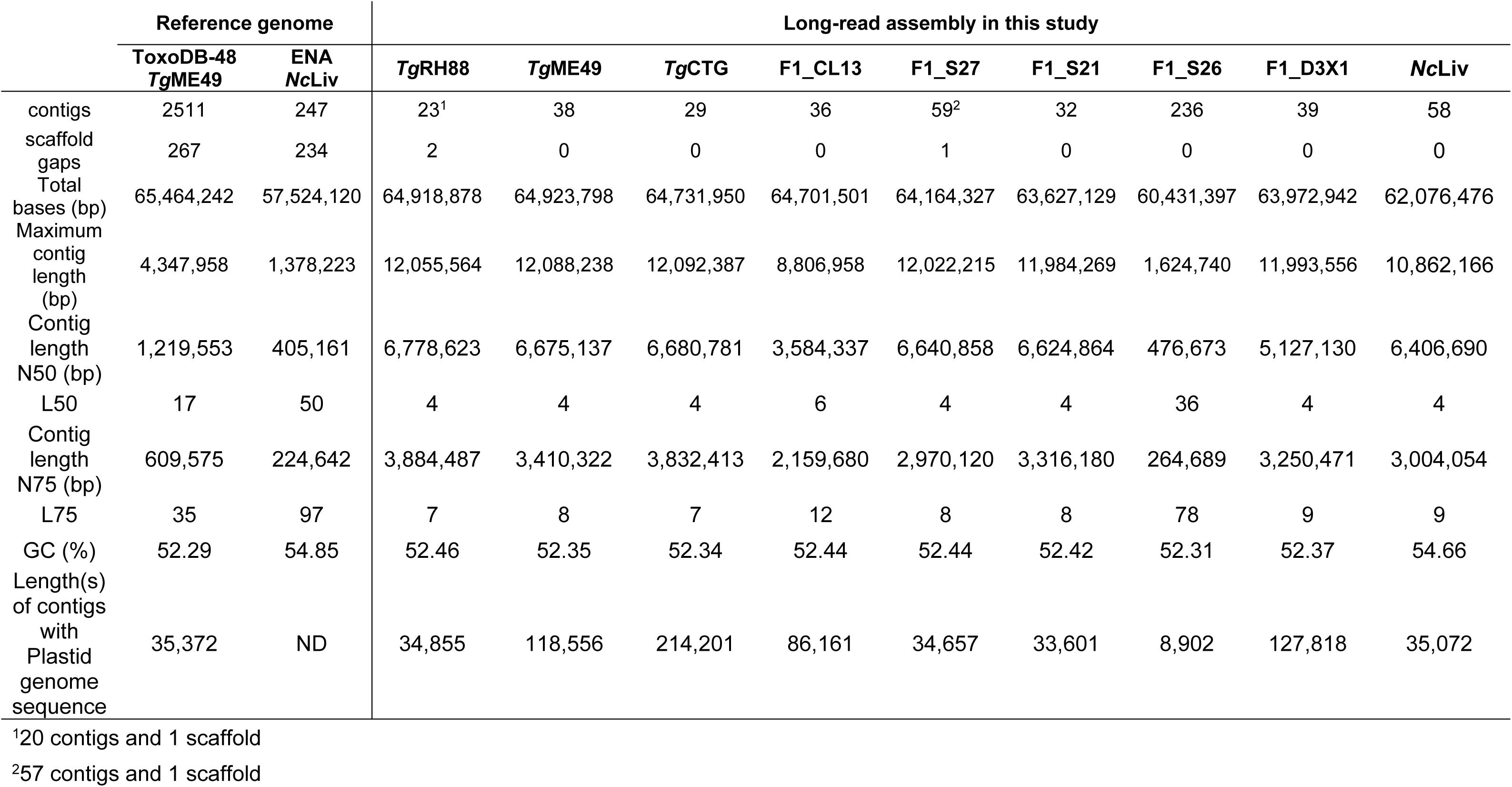
Metrics of long-read assemblies and the reference genomes after polishing.

|  | Reference genome |  | Long-read assembly in this study |  |  |  |  |  |  |  |  |
| --- | --- | --- | --- | --- | --- | --- | --- | --- | --- | --- | --- |
|  | ToxoDB-48<br>*TgME49* | ENA<br>*NcLiv* | *TgRH88* | *TgME49* | *TgCTG* | F1_CL13 | F1_S27 | F1_S21 | F1_S26 | F1_D3X1 | *NcLiv* |
| contigs | 2511 | 247 | 23<sup>1</sup> | 38 | 29 | 36 | 59<sup>2</sup> | 32 | 236 | 39 | 58 |
| scaffold gaps | 267 | 234 | 2 | 0 | 0 | 0 | 1 | 0 | 0 | 0 | 0 |
| Total bases (bp) | 65,464,242 | 57,524,120 | 64,918,878 | 64,923,798 | 64,731,950 | 64,701,501 | 64,164,327 | 63,627,129 | 60,431,397 | 63,972,942 | 62,076,476 |
| Maximum contig length (bp) | 4,347,958 | 1,378,223 | 12,055,564 | 12,088,238 | 12,092,387 | 8,806,958 | 12,022,215 | 11,984,269 | 1,624,740 | 11,993,556 | 10,862,166 |
| Contig length N50 (bp) | 1,219,553 | 405,161 | 6,778,623 | 6,675,137 | 6,680,781 | 3,584,337 | 6,640,858 | 6,624,864 | 476,673 | 5,127,130 | 6,406,690 |
| L50 | 17 | 50 | 4 | 4 | 4 | 6 | 4 | 4 | 36 | 4 | 4 |
| Contig length N75 (bp) | 609,575 | 224,642 | 3,884,487 | 3,410,322 | 3,832,413 | 2,159,680 | 2,970,120 | 3,316,180 | 264,689 | 3,250,471 | 3,004,054 |
| L75 | 35 | 97 | 7 | 8 | 7 | 12 | 8 | 8 | 78 | 9 | 9 |
| GC (%) | 52.29 | 54.85 | 52.46 | 52.35 | 52.34 | 52.44 | 52.44 | 52.42 | 52.31 | 52.37 | 54.66 |
| Length(s) of contigs with Plastid genome sequence | 35,372 | ND | 34,855 | 118,556 | 214,201 | 86,161 | 34,657 | 33,601 | 8,902 | 127,818 | 35,072 |

<sup>1</sup>20 contigs and 1 scaffold<sup>2</sup>57 contigs and 1 scaffold

The final assemblies of *Tg*RH88, *Tg*ME49, or *Tg*CTG consisted of 13 chromosome contigs/scaffolds and varying numbers of unplaced fragments, with an average total size of ∼64.8 Mb (**Table 2**). The polished *N. caninum* Liverpool assembly was composed of 58 contigs, showing a cumulative size of 62.1 Mb (**Table 2**). For all species and strains, the long reads and high coverage led to highly improved contiguity of our assemblies compared to the reference genomes. As reported in **Table 2**, with one exception (IIxIII F1 progeny S26), all the *T. gondii* final assemblies were composed of 23-59 contigs, representing a 43-109-fold reduction in the number of contigs in comparison to the ToxoDB-48_*Tg*ME49 assembly (2511 contigs; **Table 2**). We performed one-to-one mappings between our Nanopore chromosome-sized contigs and all aligning contigs from the ToxoDB-48_TgME49 assembly (Figure S1A) using Minimap2 and found that the most significant contribution to overall genome structure was the elimination of nearly all of the breaks within the ToxoDB-48_TgME49 scaffolds that are indicated within that assembly as strings of N’s at least 100 bp long (Figure S1A). We also compared the sequences and to identify new sequence uniquely incorporated into the Nanopore chromosomes and identified 42 regions of at least 10,000 bp ranging in size from 10,176 to 49718 bp (summing to 926 kb in total; yellow boxes in Figures S1A).

For *N. caninum*, compared to the existing assembly of the *Nc*Liv genome (Reid *et al*. 2012), the total number of contigs in the *Nc*Liv final assembly was reduced from 247 to 58 (**Table 2**). The contiguity improvement of long-read assembly was also evident by high contig length N50 values and low L50 values in these assemblies (**Table 2**). Furthermore, for all the long-read assemblies, 55.2% of all chromosomes were fully assembled in single contigs and flanked by two telomeric repeats with proper orientation (e.g., one end 5’-3’ TTTAGGG and the other 5’-3’ CCCTAAA; **Table S5**). This is in contrast to version 48 of the *T. gondii* ME49 genome housed at ToxoDB where 3 of the 14 chromosome assemblies have telomeric repeats on both ends, 7 have a telomeric repeat on one end, and the remaining 4 putative chromosomes have no telomeric repeats at all. Upon aligning our assemblies to their respective reference genomes (**Table S1**), we found that over 95.9% of the *Tg*RH88, *Tg*ME49, or *Tg*CTG final assembly sequences could be mapped to their relevant reference genome. The alignments revealed that our *de novo* assemblies had fewer mismatches and indels per 100 Kb and higher longest alignment and total aligned length when compared to the assemblies housed on ToxoDB (**Table 3**). To assess genome assembly completeness, we used BUSCO analysis on the polished *Tg*RH88, *Tg*ME49, and *Tg*CTG assemblies and compared the results to a similar analysis for the unpolished assemblies. This analysis, which counts the number of single-copy orthologs unambiguously identified in a genome assembly, found that for 215 such loci 88% of the complete genes were recovered from the polished, long read based assemblies, while 23.3 - 54% were identified in the unpolished assemblies (**Table 3**). These data show the utility of our sequence polishing for improving the accuracy of the assembly.

**Table 3.** Metrics of the long-read assemblies before and after polishing.

|  | *TgRH88* |  | *TgME49* |  | *TgCTG* |  |
| --- | --- | --- | --- | --- | --- | --- |
|  | Initial<br>assembly | Final<br>assembly | Initial<br>assembly | Final<br>assembly | Initial<br>assembly | Final<br>assembly |
| **Contiguity** |  |  |  |  |  |  |
| # contigs/scaffolds | 23 | 21 | 38 | 38 | 38 | 29 |
| Total bases (bp) | 64,401,064 | 64,918,878 | 64,522,756 | 64,923,798 | 64,789,158 | 64,731,950 |
| Maximum contig length(bp) | 11,930,269 | 12,055,564 | 12,002,493 | 12,088,238 | 12,040,189 | 12,092,387 |
| Contig/scaffold length N50 (bp) | 6,718,904 | 6,778,623 | 6,635,075 | 6,675,137 | 6,653,560 | 6,680,781 |
| **Accuracy** |  |  |  |  |  |  |
| Genome fraction (%) | 97.514 | 97.470 | 95.914 | 95.915 | 98.367 | 98.353 |
| # mismatches per 100 Kbp | 85.91 | 57.33 | 61.34 | 49.82 | 47.85 | 44.11 |
| # indels per 100 Kbp | 706.9 | 35.1 | 537.5 | 26.59 | 348.43 | 21.94 |
| Largest alignment (bp) | 2,737,008 | 2,768,614 | 4,425,732 | 4,452,657 | 2,573,606 | 2,584,520 |
| Total aligned length (bp) | 62,839,196 | 63,538,670 | 63,828,373 | 64,235,943 | 63,473,895 | 63,611,317 |
| **Completeness** |  |  |  |  |  |  |
| Complete BUSCOs protists (%) | 23.3 | 88.9 | 39.1 | 91.6 | 54.0 | 91.2 |
| Fragmented BUSCOs protists (%) | 0.9 | 0.0 | 1.4 | 0.0 | 1.9 | 0.0 |
| Missing BUSCOs protists (%) | 75.8 | 11.1 | 59.5 | 8.4 | 44.1 | 8.8 |

### Long-read assembly identifies 13 chromosomes in multiple *T. gondii* genomes

To assess the structural correctness of all of the the long-read assemblies, we aligned our *T. gondii* assembly sequences to the reference genome using Nucmer. As can be seen in **Figure S1B**, all of the *T. gondii* long-read assemblies exhibited strong collinearity with their corresponding reference genomes, barring a small number of putative inversions. Consistent with our finding for *Tg*RH88 primary assembly, the chr VIIb/VIII fusion was observed in each of our *T. gondii* assemblies (*Tg*ME49 is represented in **Figure 3A**, red box; and **Figure S1A,B**). To further confirm this observation, we aligned our *Tg*ME49 corrected reads back against the *Tg*ME49 long-read assembly and found an average read depth of 40× for the entirety of TgME49_tig00000001_chrVIII, and 37× for the “breakpoint” (TgME49_tig00000001_chrVIII: 5090422 bp) of chrVIIb and chrVIII, indicating that this was unlikely due to an assembly error (**Figure 3B**). We then mapped the corrected nanopore reads again to the ToxoDB-48_*Tg*ME49 reference genome, and the alignments showed that all of the reads that were mapped either to the end of chrVIIb or to the beginning of chrVIII spanned the gap between the two chromosomes, with an average coverage of 105× (**Figure 3C**). This link between reads aligning to chromosome ends was not present in any other chromosome pair (for instance, between chromosomes IX and X; **Figure 3D**).

**Figure 3.**
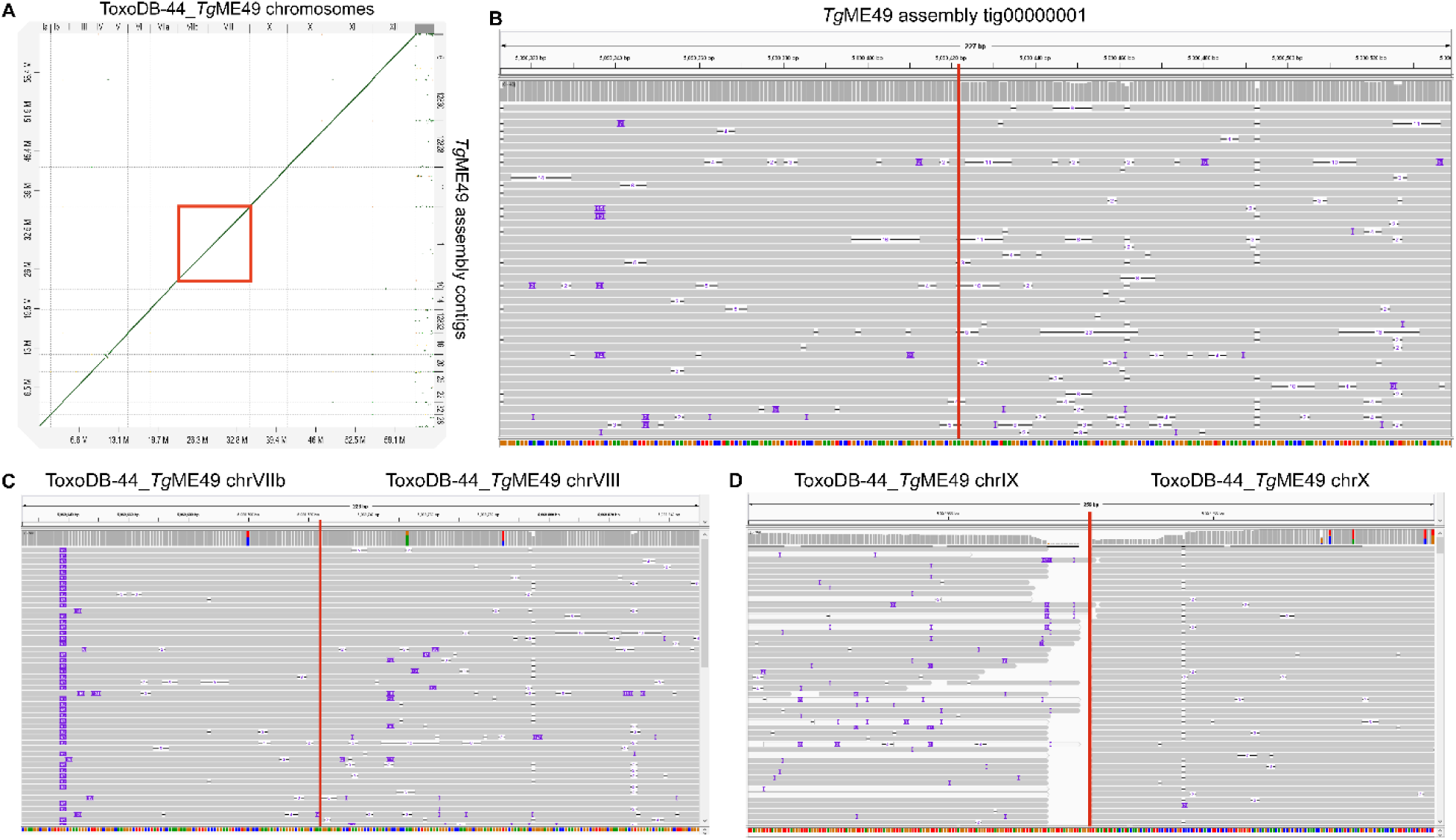
Long-read assembly identifies 13 chromosomes in the T. gondii genome from multiple strains. **(A)** Dot plot showing the comparison of the TgME49 long-read assembly and the ToxoDB-48_TgME49 genome. Red box shows that the chromosomes VIIb and VIII in the ToxoDB-48_TgME49 genome are fused in a single contig, TgME49_tig00000001_chrVIII, in the TgME49 long read assembly. (B) Coverage of the “breakpoint” (TgME49_tig00000001_chrVIII: 5090422 bp, indicated by a vertical red line) of chromosomes VIIb and VIII with 37 Nanopore reads in the TgME49 long-read assembly. (C) Coverage of the edges (indicated by a vertical red line) of chromosomes VIIb and VIII with 105 Nanopore reads mapped to the ToxoDB-48_TgME49 genome. (D) Nanopore reads mapping to the end of chromosomes IX and X in the ToxoDB-48-TgME49 genome assembly, showing that Nanopore reads only map to the end of each chromosome and do not span the junction between these chromosomes (indicated by a vertical red line).

This observation of a fusion between chrVIIb and chrVIII was in agreement with the observations in our TgRH88 assembly described above (**Figure 1C-E**). Similarly, when we mapped Hi-C data from (Bunnik *et al*. 2019) was aligned to our *Tg*ME49 long read assembly, the resulting interchromosomal contact-count map revealed 13 chromosomes in the *Tg*ME49 long read assembly by showing that each chromosome exhibited a single centromeric interaction with each other chromosome, and that TgME49_tig00000001_chrVIII was a complete single chromosome (**Figure S2**). The intrachromosomal contact-count map of TgME49_tig00000001_chrVIII showed a strong and broad diagonal and no aberrant signal along the contig or at the the “breakpoint” of chrVIIb and chrVIII (**Figure S2B**). Similar patterns were also observed in our *Tg*CTG, S27, and S21 assemblies (**Figure S2B**). Collectively, these data show the *T. gondii* karyotype has been incorrectly calculated and contains 13, rather than 14, chromosomes. We refer to this fused chromosome as chrVIII in our assembly and have eliminated chrVIIb.

The *Tg*CTG assembly had chromosome scale resolution, where 13 contiguous sequences (contigs) corresponded to the 13 chromosomes. However, chromosome VIIa in the *Tg*ME49 assembly and chromosome XI in the TgRH88 assembly was spread across two contigs. Hi-C data has historically been used to improve genome assemblies on the basis of contact frequency depending strongly on one-dimensional distance (Dudchenko *et al*. 2017). That is, Hi-C alignment to contigs in the correct order and orientation would reveal the canonical intrachromosomal pattern of enriched interactions along the diagonal (where one-dimensional genomic distance between bins is the smallest). Hi-C alignment to contigs in the incorrect order and orientation would be illustrated by patterns associated with large-scale inversions, where there is a high interaction frequency between bins placed far away from each other. Leveraging this, we were able to determine the order and orientation of the contigs for chromosomes ChrVIIa and XI in ME49 and RH88, respectively. These changes were incorporated prior to submission of the assemblies to Genbank.

### Long-read assembly detects structural rearrangements in the *T. gondii* genome

In addition to repeated elements, our assembly was capable of detecting gross structural rearrangements (insertions, deletions, inversions, relocations, and translocations) since the assembly based on long reads was highly contiguous and had near chromosome scale resolution. We aligned the final assemblies of the 8 *T. gondii* strains to the reference genomes and searched for structural variants using MUMmer (**Figure S1B**). Consistent with the data shown in **Figure S1B**, most contigs in the long-read assemblies were collinear with the chromosomes in the ToxoDB-48 genomes, but not always in a 1:1 correspondence. Interestingly, we observed a 15.7 Kb inversion on chrIII in our *Tg*RH88 assembly, which was absent in *Tg*ME49, *Tg*CTG, or any F1 progeny assembly (**Figure 4A**). We also detected another ∼20 Kb inversion on chrXII, which was present in *Tg*ME49, *Tg*CTG, and the F1 progeny assembly, but not in the *Tg*RH88 assembly (*Tg*ME49 is represented in **Figure 4B**).

**Figure 4.**
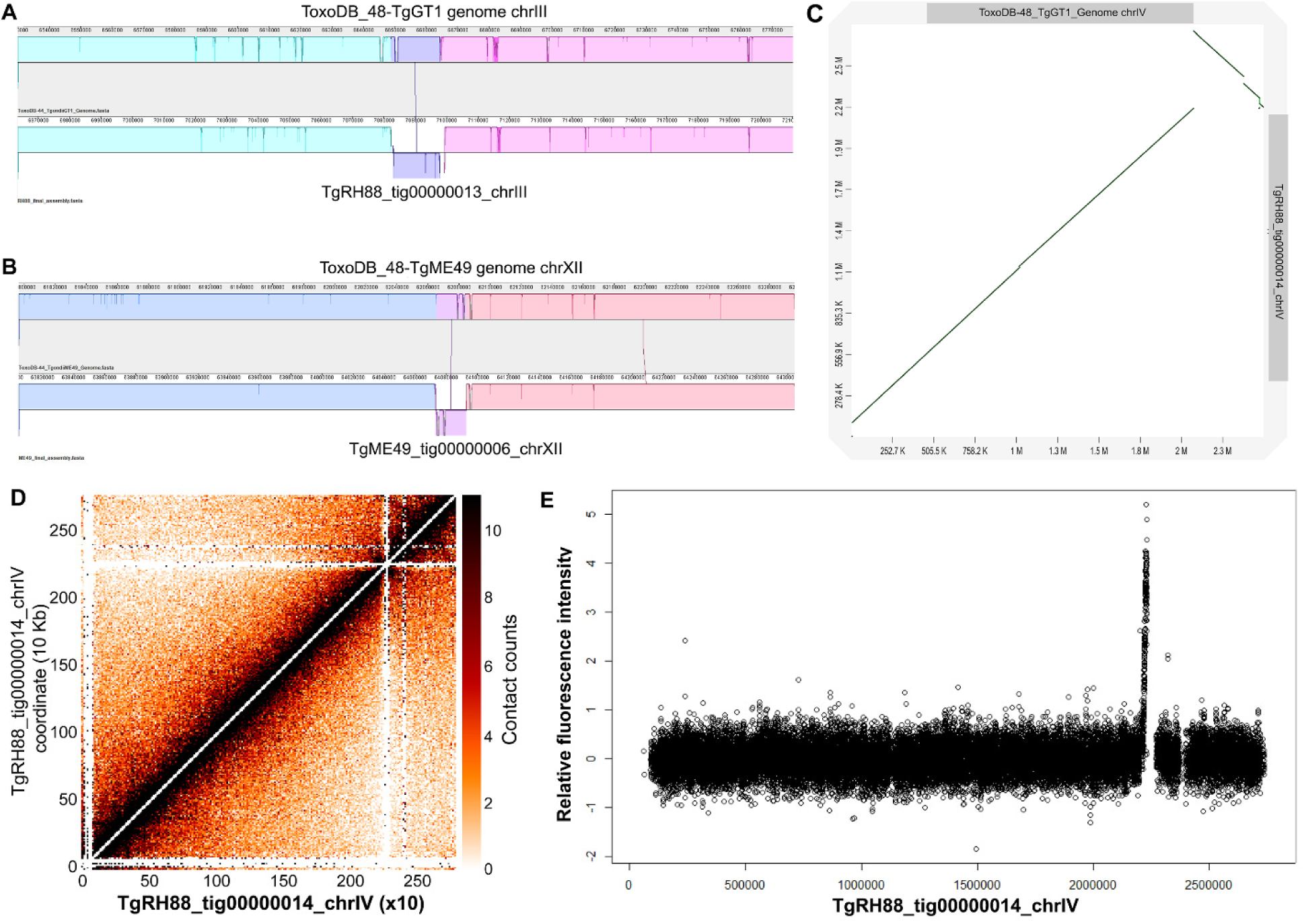
Long-read assembly reveals previously unknown inversions and the centromere location on chrIV in T. gondii. (A) Inversion in the RH88 long-read assembly on chromosome III relative to the ToxoDB-48-TgGT1 assembly. (B) Inversion in the ME49 long-read assembly on chromosome XII relative to the ToxoDB_44-TgME49 genome. (C) Dot plot comparison of the TgRH88 long-read assembly and the ToxoDB-48_TgGT1 genome showing a 429.3-Kb inversion at 2,096,529-2,525,795 bp on chrIV. (D) Intrachromosomal Hi-C contact-count heat map plotted using the sequence of contig_14 in TgRH88 long-read assembly showing a clear centromere signal at position 2.2-2.3 Mb. (E) ChIP-on-chip signal of centromeric histone 3 variant (CenH3) (Brooks et al. 2011) plotted using the TgRH88 long-read assembly as coordinate.

Centromeres for 12 of the 13 *T. gondii* chromosomes have been identified using ChIP-on-chip (chromatin immunoprecipitation coupled with DNA microarrays) of centromeric and peri-centromeric proteins, but locations of centromere of chrVIIb and chrIV remain unknown (Brooks *et al*. 2011; Gissot *et al*. 2012). For chrVIIb, no hybridization of the centromeric probe to the genomic chip was detected in the ChIP-on-chip assay (Brooks *et al*. 2011), which could be explained by our observation that chrVIIb and chrVIII are a single chromosome, and the centromere of this large chromosome appears to be in the center of this “fused” chromosome, in an area that was previously thought to be the beginning of chromosome VIII (**Figure 1E**). For chromosome IV, two inconsecutive peaks of hybridization were detected at positions 2,501,171-2,527,417 bp on chrIV and 1-9968 bp in the unplaced contig AAQM03000753 in the *Tg*GT1 genome based on published ChIP-on-chip data (Brooks *et al*. 2011). Our *Tg*RH88 assembly successfully relocated the sequences in AAQM03000753 into chrIV and revealed a 430.9-Kb inversion event at 2,096,529-2,527,423 bp on chrIV relative to the ToxoDB GT1 reference genome (**Figure 4B**). This inversion was unlikely to be due to an assembly error since it was shown in all of our *T. gondii* assemblies, and was supported by alignment of 128 canu-corrected reads spanning the boundaries of the *Tg*RH88 assembly. When we mapped the ChIP-on-chip data obtained from (Brooks *et al*. 2011) to our *Tg*RH88 assembly (after remapping the probe sequences to our TgRH88 assembly), we resolved the chrIV centromere to a single significant signal peak at 2.20-2.23 Mb on chrIV (**Figure 4C** and **Figure S3**). This finding was also supported by published Hi-C data re-aligned to our Nanopore assemblies, since the intrachromosomal contact count map showed a clear interchromosomal contact signal at 2.20-2.30 Mb on chrIV in the *Tg*RH88 assembly (**Figure 4B**). Collectively, our data not only resolved the molecular karyotype of *T. gondii*, but also resolved the precise location of the chrIV centromere.

### Long-read assembly adds new sequences to the *T. gondii* reference genome

As shown in **Figure 1C** and **Table S5**, each chromosome-sized contig of our long-read *de novo* assemblies was longer than its cognate chromosome of the ToxoDB-48 *T. gondii* reference genome, and for TgRH88, TgME49 and TgCTG we were able to assemble between 1.7 and 3.9 Mb of previously unlocated and/or unassembled sequence to the chromosomes of these assemblies. These new sequences were scattered across the genome, and filled in nearly all of the sequence gaps found in the reference genome. The new sequences added by the long-read assemblies extended the subtelomeric regions of the chromosomes in the *T. gondii* reference genome. While 4 chromosomes of the ToxoDB-48_*Tg*ME49 genome contained no telomeric repeat and 7 were missing one of the telomeric repeats, all of the chromosome contigs in the *Tg*ME49 long-read assembly were assembled up until both telomeric caps (**Table S5**). Both telomeres were found in 12 out of the 13 chromosomes in the *Tg*CTG long-read assembly, and one chromosome contig lacked one of the telomeric repeats, whereas only 5 chromosomes in the ToxoDB-48_*Tg*VEG genome contained one telomere and no telomeric repeat was found in the rest of the chromosomes (**Table S5**). Similarly, both telomeres in 7 chromosomes and one telomere in 6 chromosomes were resolved in the *Tg*RH88 long-read assembly, while there were only 2 chromosomes in ToxoDB-48_*Tg*GT1 genome that contained one telomere (**Table S5**).

In addition to telomeres, the bulk of the remaining new sequence was due to tandem arrays of sequence (both coding and non-coding). For example, two repetitive gene sequences are used for high sensitivity detection of *T. gondii* in tissue and environmental samples, the B1 gene (Burg *et al*. 1989) and the so-called “529 bp repeat” (Reischl *et al*. 2003; Edvinsson *et al*. 2006). The precise copy number for these genes has been impossible to determine using 1st and 2nd generation sequencing technologies and/or molecular biological experiments like Southern Blotting. Therefore, we used a curated blastn approach to quantify copy number for each of these sequences across our respective Nanopore assemblies. As shown in **Figure 5A**, copy number for the B1 gene was dramatically higher in our Nanopore assemblies compared to existing ToxoDB assemblies (as expected). What was unexpected, however, was that copy number for this gene was lower than that predicted in the literature, ranging between 9 and 19 tandem copies depending on the strain (**Figure 6A**), compared to quantitative blotting-based estimates of 35 (Burg *et al*. 1989). Copy number at this locus was stable, in that for all of the queried IIxIII F1 progeny copy number for each was identical to the parent from which it obtained that chromosomal segment. In contrast to the B1 locus, the “529 bp repeat” locus varied dramatically between isolates and these same F1 progeny. Copy number ranged from 85 to 205, and copy number at this locus for all F1 progeny varied independently of the underlying genotype for that region (white letters and green/blue in **Figure 5B**). The size of this genome expansion is best illustrated by the whole chromosome alignment shown in **Figure 5E** comparing the *T. gondii* ME49 529 bp repeat locus in the version 48 assembly on ToxoDB to our Nanopore assembly (**Figure 5E**). Importantly the 529 bp repeat locus occurs near a sequence assembly gap, and our long read assembly closed this gap (see below and **Figure 5D**), giving the most accurate estimate of 529 bp repeat copy number in any *T. gondii* strain which again varies compared to estimates in the literature (ranging from 200-300 copies; e.g., (Reischl *et al*. 2003; Edvinsson *et al*. 2006)). Regardless, similar to tandem gene arrays discussed above in **Figure 6**, it appears that even noncoding repeats like the B1 gene and the 529 bp repeat can also change in number irrespective of whether there is sexual recombination. While the 529 bp repeat and B1 gene are found in a single locus, other tandem repeats like TgIRE and SAT350 are found across multiple subtelomeric genomic locations (Echeverria *et al*. 2000; Clemente *et al*. 2004). We mapped these sequences across the genome in all of our sequenced *T. gondii* strains, and again found different results depending on the queried locus. For the TgIRE sequence, a 1919 bp repeat, chromosome-wide copy number in our sequenced strains was ∼2x that found in the reference sequences (Figure 6C), but overall copy number was similar across all of our Nanopore-derived sequences. The SAT350 sequence was also found at much higher copy number in our Nanopore assembled chromosomes compared to the reference sequences (Figure 6D), but was more variable in the F1 progeny clones. This could be due to changes during sexual recombination as for the 529 bp repeat above, or differences in sequence coverage for the F1 progeny. Regardless, our approach has provided better resolution of copy number for 4 well-known tandem repeat sequences, and allowed us to determine which may be more susceptible to copy number change. Moreover, unappreciated strain differences in copy number at these loci may adversely affect interpretation of PCR-based detection assays, especially those using more quantitative methods.

**Figure 5:**
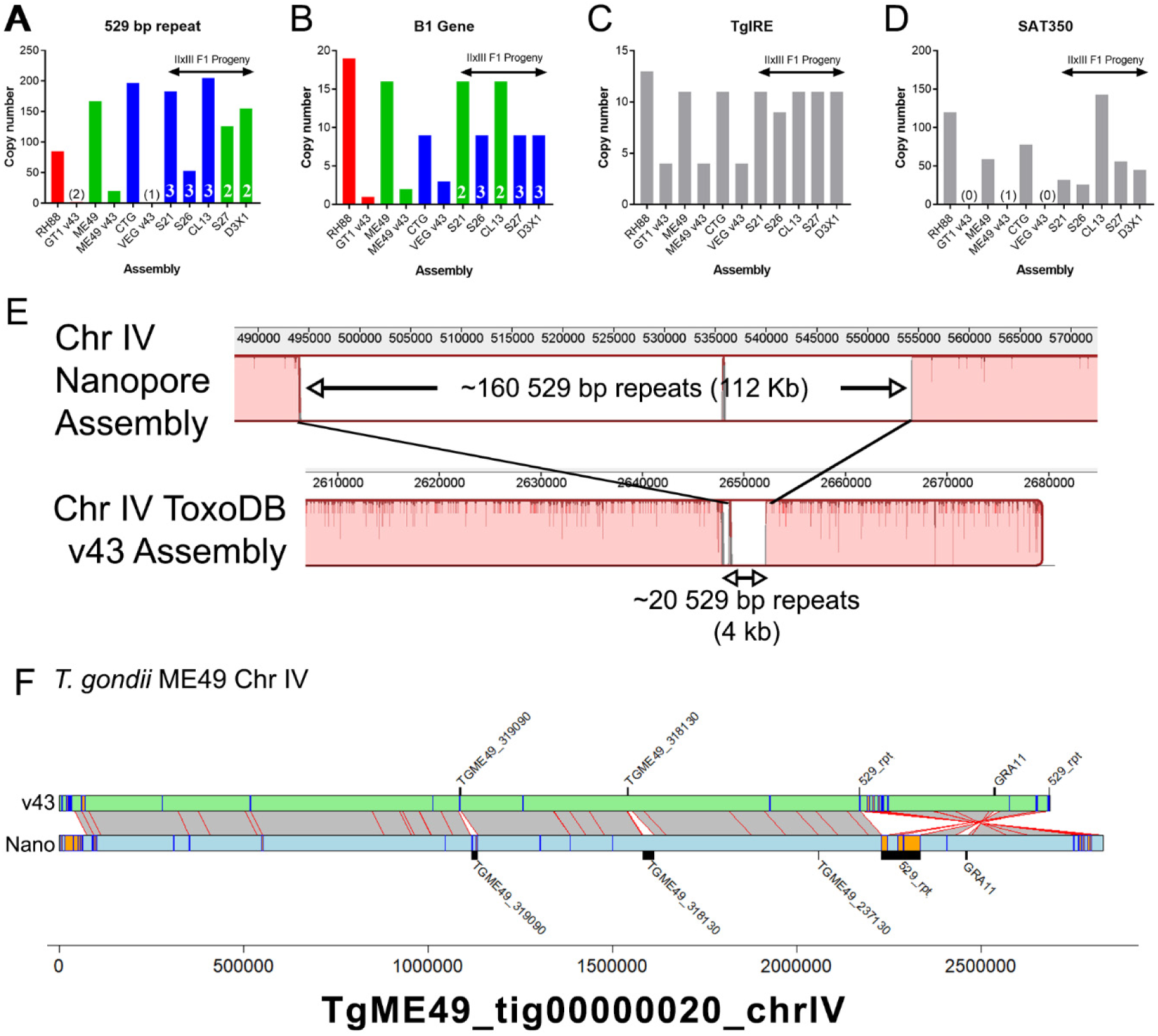
Long read sequence assemblies precisely resolve canonical repeat sequences and identify additional expansions at gene-harboring loci. (A-D) Estimated copy number for Nanopore assemblies and existing genome assemblies on ToxoDB (“v48”) for T. gondii strain types 1, 2 and 3 and IIxIII F1 progeny. In all cases, Nanopore assemblies identified higher numbers of each repeat locus. In the F1 progeny, B1 gene copy number tracked directly with the genotype (type 2 or 3) at that locus (A), while these same F1 progeny harbored unique numbers of 529 bp copies, all of which were not only distinct from their respective genotypes of origin but distinct from one another (B). The TgIRE and SAT350 repeats also were better resolved in our Nanopore assemblies (C,D) although determining genotype of the corresponding region is not possible since these repeats are found at multiple locations throughout the genome. (E) Whole chromosome alignment focused on the 529 bp repeat region for the v48 ToxoDB assembly (bottom) and our Nanopore-based assembly (top). Expansion of the known genome sequence at this locus in the Nanopore sequence compared to the ToxoDB assembly is clear, and consistent with our identification of ∼140 previously unknown 529 bp repeats in the ME49 genome. (F) Alignment and annotation of repeat sequences of ME49 v48 ToxoDB chromosome IV and that from our polished Nanopore assembly. Grey bars with red borders indicate mapping regions ≥10,000 bp determined using nucmer, while orange boxes with blue borders indicate tandem repeats with period sizes ≥ 500 bp and at least 2 copies. Bars that appear orange are larger than those that are only blue. Incorrect inversion on the right arm of chromosome IV in the ToxoDB assembly is evident, as is the more accurately resolve 529 bp repeat locus that was likely a cause for the inversion in standard assemblies from multiple strains.

**Figure 6.**
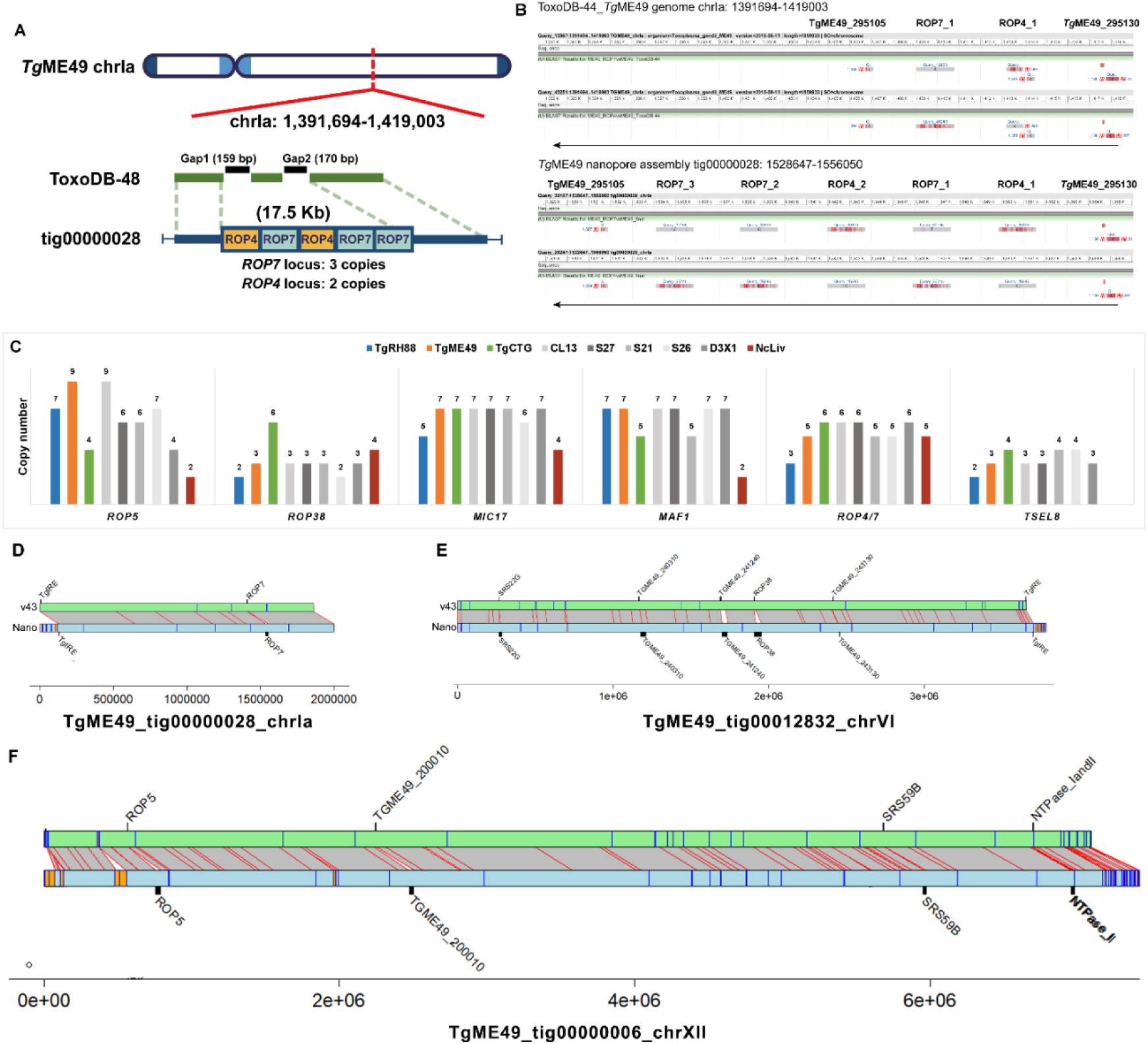
Long-read assembly resolves duplicated locus structure in T. gondii genome. (A) Two unresolved scaffold gaps on chrIa in ToxoDB-48_TgME49 genome span a 17.5-Kb tandem repeat containing multiple copies of ROP4 and ROP7. The ROP4/7 gaps are closed by the TgME49 long-read assembly (TgME49_tig00000028), revealing a tandem array of 5 copies of this gene in the order shown. (B) BLASTN alignment of the ROP4/ROP7 coding sequence in the ToxoDB-48_TgME49 genome (upper panel) and the TgME49 long-read assembly (lower panel). (C) Copy number determination at 6 canonical tandem gene arrays across 8 T. gondii strains and 1 N. caninum strain. Data from CL13, S27, S21 and S26 show that copy number can change during sexual recombination since copy number in these F1 progeny clones do not match copy number in either parent. (D-F) Whole chromosome alignments between ME49ToxoDB-48 and our Nanopore assemblies at loci harboring tandem gene arrays. Grey boxes with red borders indicate 1-to-1 mapping regions ≥ 10,000 bp determined by nucmer and orange/blue boxes are as described in Figure 5. Black bars indicate size of select tandem repeats in the ToxoDB and Nanopore assemblies.

With respect to chromosome IV, mapping the 529 bp repeat to our Nanopore assemblies while comparing them to reference genomes in ToxoDB identified what appears to be an inversion in the right arm of chromosome IV (Figure 6F). This inversion was present in the *T. gondii* VEG and GT1 reference genomes, suggesting that it was due to systematic assembly error. This inversion in the ToxoDB reference genome is flanked by multiple tandem repeats (identified using tandem repeats finder; blue/orange boxes in **Figure 6F**). In addition, the 529 bp repeat can be found at the end of each inversion (“529_rpt”, **Figure 6F**) in the ToxoDB chromosome, but in our Nanopore assembly the 529 repeat cluster is only found in one location (where it is greatly expanded compared to those in the ToxoDB reference). These data provide strong evidence that chromosome IV is incorrectly assembled in multiple ToxoDB reference genomes due to misassembly of the 529 bp repeat locus, and our Nanopore assembly has resolved this discrepancy in multiple *T. gondii* genomes.

### Long-read assembly resolves tandem duplicated locus structure in the *T. gondii* genome

Many scaffold gaps within chromosomes remain unresolved (**Table 2**) in the *T. gondii* reference genome particularly in the regions that contain repeated elements. Our long read assembly closed nearly all of the gaps in the *T. gondii* and *N. caninum* genomes (Tables 2 and S6). Furthermore, many unplaced sequences in the *T. gondii* reference genome were assembled into contigs in our assembly. For instance, the unplaced contig KE140372 in the ToxoDB-48_*Tg*ME49 genome, which was 2194 bp in length and contained a sequence encoding a rhoptry protein 4 paralog, was assembled in TgME49_tig00000028_chrIa in our *Tg*ME49 assembly. Unplaced contigs in the ToxoDB-48_*Tg*GT1 genome, AAQM03000823 and AAQM03000824, were assembled in TgRH88_tig00000013_chrIII in our *Tg*RH88 assembly.

We have had a long-standing interest in variation at tandemly expanded gene clusters and how this affects *T. gondii* virulence which was first spurred by our identification of ROP5 gene cluster as being a critical determinant of virulence in the mouse (Reese *et al*. 2011). Our genome-wide analyses of copy number variation across multiple strains and species (Adomako-Ankomah *et al*. 2014) identified 53 putative tandemly expanded gene clusters in *T. gondii* (shown in Table S6), some of which (such as MAF1) have been shown to be important in host pathogen interactions (Adomako-Ankomah *et al*. 2016). In that initial study we used sequence coverage to infer copy number and we were eager to refine this analysis using the long-read de novo assemblies presented here.

The gap closure enabled us to determine the number, order and orientation of *T. gondii* duplicated loci including *ROP7*, *ROP5*, *ROP38*, *MIC17*, *MAF1*, and *TSEL8*. The *ROP7* locus is represented in **Figure 6A**, where we identified two unresolved scaffold gaps on chrIa in the ToxoDB-48_*Tg*ME49 genome (black bars), and these gaps marked the site of *ROP4/7* locus. This entire region was spanned by a single contig, TgME49_tig00000028_chrIa, in our *Tg*ME49 assembly (the blue bar in **Figure 6A**). Aligning the *ROP7* genomic sequence (ToxoDB: TGME49_295110) against TgME49_tig00000028_chrIa using BLASTN revealed 3 copies of the *ROP7* repeat, while only one was predicted in the ToxoDB-48_*Tg*ME49 genome (**Figure 6A and 6B**) Interestingly, while the ToxoDB-48_*Tg*ME49 genome identified one copy of the *ROP4* gene (GenBank: EU047558.1), our *Tg*ME49 assembly showed that 2 copies of *ROP4* exist in this locus, one of which was found between the first and the second copy of *ROP7* (**Figure 6B**). To validate this finding, we identified 12 individual Canu-corrected reads that spanned this entire tandem array, and each one that we examined provided evidence for 3 copies of *ROP7* and 2 copies of *ROP4* arranged in the order ROP4_1-ROP7_1-ROP4_2-ROP7_2-ROP7_3 (**Table S6**). The copy number and copy order of other known tandem locus expansions (taken from the supplementary table in (Adomako-Ankomah *et al*. 2014)) in the strains we sequenced were identified and are listed in **Table S6**. Surprisingly, we observed changes in copy number at some of these loci when we compared the parental (*Tg*ME49 or *Tg*CTG) and the progeny (CL13, S27, S21, S26, and D3X1) assemblies (**Figure 6C**). For example, while the *ROP5* locus had 9 copies in our *Tg*ME49 assembly and 4 in our *Tg*CTG assembly, it harbored 6 copies in the F1 progeny S27 and S21, and 7 in S26 (**Figure 6C**). There was an array of 7 tandem copies of *MIC17* in *Tg*ME49 and *Tg*CTG assemblies, whereas it was present in 6 copies in the S26 assembly (**Figure 6C**). These data indicated that changes in copy number and order at tandem gene arrays can occur during sexual recombination. Moreover the ease with which a Nanopore assembly can be generated and assembled provides a new way to assess the occurrence and impact of acute changes in copy number that occur during asexual and sexual propagation of parasites like *T. gondii*. Whole chromosome alignments are shown for chromosomes Ia, VI and XII to further illustrate the increase in sequence size at the chromosome level for loci like ROP7, ROP38 and ROP5, as well as other known tandem gene arrays (**Figure 6D, E and F**).

We performed a similar analysis genome-wide for the remaining previously identified tandem expansions ((Adomako-Ankomah *et al*. 2014); **Table S7**), and were able to further curate this list of putative repetitive loci. While in some cases our predictions from shotgun sequence read coverage was similar to that predicted using Nanopore assembly (as for those genes described in **Figure 6** as well as many others in **Table S7**), for some we were able to determine that there was in fact no evidence for the presence of a tandem gene array at that locus. For example expanded locus 9 (EL9; TGME49_319090), EL19 (TGME49_204560) and EL27 (TGME49_264420) are all likely to be single copy genes based on our Nanopore assemblies (**Table S7**), even though based on sequence coverage alone they ranged in predicted copy number between 15 and 60 (**Table S7** and (Adomako-Ankomah *et al*. 2014)). While the reasons for each of these discrepancies will be locus specific, it is likely that the overestimation of copy number using sequence coverage in our prior study was due to the presence of low complexity sequence (e.g., short tandem repeats) across the single copy gene rather than being due to actual tandem expansion. Overall, our current assemblies refine and further curate this locus list, providing the most accurate estimate to date of which loci encode tandem gene arrays and how their copy number varies across strain and species.

As described above all of our sequence assemblies increased the size of the contiguous assemblies by 1-3 Mb. While some of this new sequence is most certainly derived from gene-poor regions containing simple tandem repeats (and some of these regions are annotated as yellow boxes in **Figure S2**), the relatively high gene density of the *T. gondii* genome led us to hypothesize that some of this “new” sequence should be derived from gene sequences that were previously masked by assembly gaps. Therefore we used BLASTN and custom parsing scripts to identify genome expansions in our Nanopore assembly relative to the v48 sequence on ToxoDB specifically using all available predicted genes as query sequences. Overall, we identified 62 gene-containing loci that were at least 10 Kb larger in our Nanopore assembly compared to ToxoDB v48, representing 1.2 Mb of sequence. These expansions are represented in **Figure S4** as yellow blocks, and are shown along with known tandem gene arrays (red blocks) and existing sequence gaps (black lines). Well known tandem gene arrays that are collapsed in 1st and 2nd generation sequence-based assemblies like MAF1 and ROP5 were identified in this analysis (**Figure S4**), confirming the accuracy of the approach. What was unexpected was the unequal distribution of these “expansions” in our genome-wide analysis (e.g., compare chromosomes XI and X). These expansions are due to increased copy number as well as previously unknown insertions of repetitive sequence. In addition to these gene-containing loci, we have identified all tandem repeats with a period size > 500 bp in our de novo assembly of *T. gondii* ME49 and estimated their copy number in our de novo assemblies of *T. gondii* RH88 and CTG (Table S8).

### Standard error correction methods for tandem gene arrays fail to remove extensive homopolymeric repeats that lead to artifactual pseudogenization

Error correction using Pilon is a common practice after generation of single molecule long-read assemblies since the overlap-based correction employed by assemblers like Canu fails to correct systematic errors in the data. For Nanopore, homopolymer runs as short as 3 bp are often truncated in the consensus [need citation], leading to extensive artificial gene pseudogenization. As described above hybrid assembly or error correction can eliminate many of these systematic errors leading to improved protein coding gene annotations (Table 3). To examine this is greater detail we used TBLASTN to align *T. gondii* ME49 (version 48) protein sequences for all single exon genes to the ToxoDB reference genome, the primary Canu assembly, and to our hybrid-corrected assembly. As shown in Figure 2A the raw Canu assembly had only 4 full length, identical sequence hits to single exon genes, while the Pilon-corrected assembly predicted nearly all of the query single exon genes with 100% identity and 100% sequence coverage (1054/1121; 94%; Figure 7A). The effectiveness of Pilon error correction is shown for 2 single exon, single copy genes (ROP16 and ROP18; Figure 8B), where the coding sequence had numerous frameshifts causing fragmented mapping in the Raw assembly but resolved to a single gene with 100% identity and sequence coverage after Pilon correction. In contrast to these single copy genes, we found that Pilon-based error correction performed much more poorly at multicopy loci like those encoding ROP5 and ROP38. As shown in **Figure 7C and D**, many of the predicted ROP38 and ROP5 coding sequences are still highly fragmented after Pilon error correction, presumably due to an inability of Pilon to assign enough reads to each copy to correct what are mostly homopolymeric repeat errors. We wrote a custom Perl script to correct remaining homopolymer errors in tandem gene arrays (using Illumina sequence read alignments to either extend homopolymers up to 10 bp or truncate by a single base pair) and found that this eliminated many of the artefactual pseudogenes for the tandem gene expansions at the ROP5 and ROP38 loci (see bottom of **Figure 7C and D**). Note that unlike single copy genes where Canu plus Pilon correction was sufficient to correct them (**Figure 7B**), we only eliminated these likely artefactual pseudogenes after running our supplemental correction scripts (see Materials and Methods for specific details about the correction script). We have deposited these corrected sequences in Genbank separately from our genome assemblies for all the loci characterized in Figure 6C (submission numbers pending).

**Figure 7:**
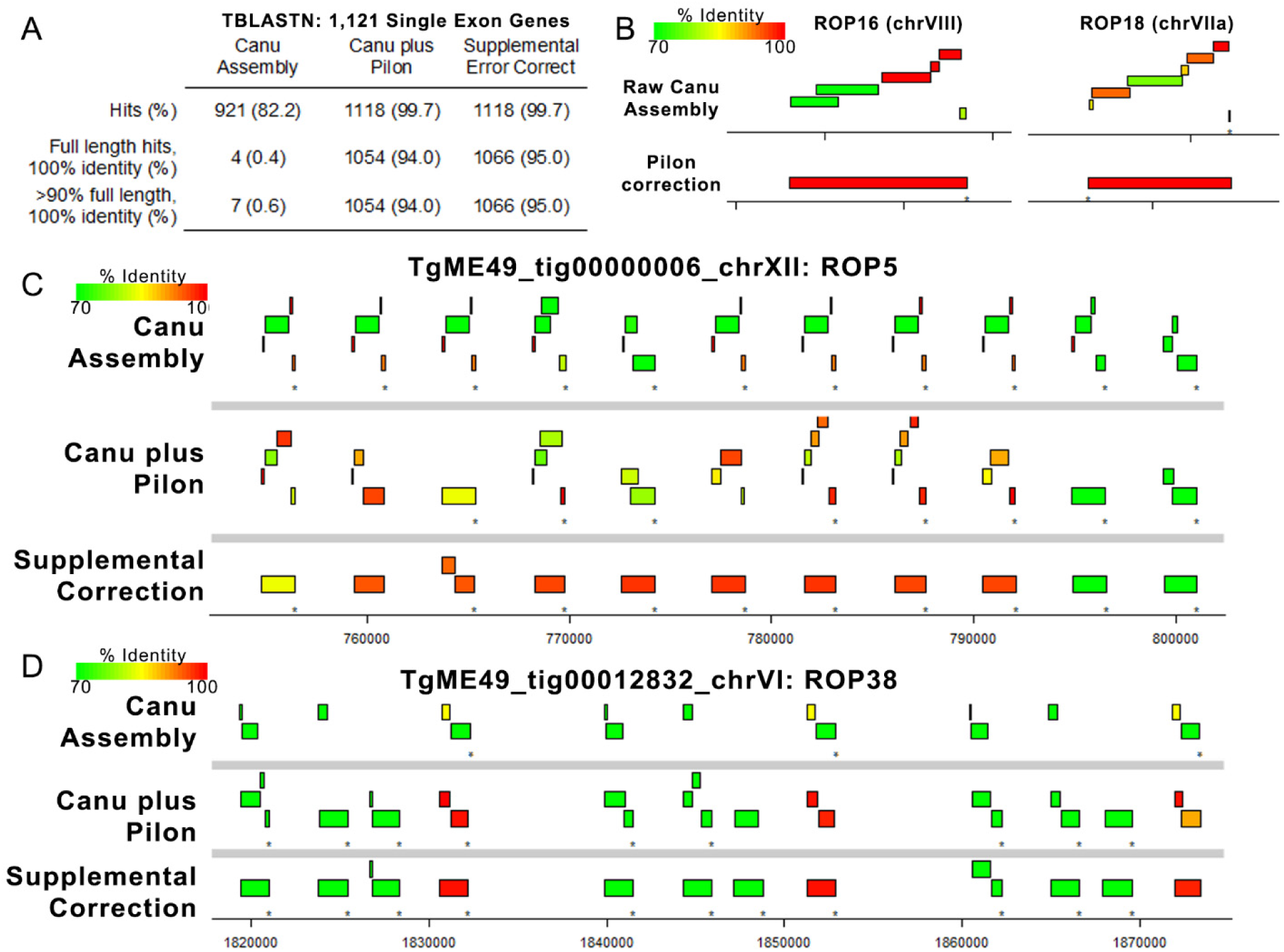
Error polishing with Pilon is effective for most single copy genes but resolution of tandem gene expansion errors requires supplemental correction. (A) We identified 1121 single exon genes with predicted proteins that mapped with 100% identity and 100% coverage using TBLASTN against the ToxoDB-48 ME49 genome. Then, we mapped these against the raw assembly generated by Canu, the polished assembly generated by 4 rounds of Pilon, and after 4 rounds of supplemental error correction. Pilon error correction was sufficient for perfect mapping of 94% of the query single exon genes (compared to only 0.4% for the raw Canu assembly), and supplemental error correction only increased this mapping percentage slightly. (B) Plots representing TBLASTN analysis of protein sequences from two single copy genes showing the improved mapping achieved by Pilon-based error correction. Mapping identity is indicated by the color of the box representing the alignment. (C-D) Plots representing protein sequence from the ROP5A (C) or ROP38 (D) gene mapped using TBLASTN against the raw Canu-only assembly, the Pilon-corrected assembly and the region corrected using our supplemental approach tailored to tandem gene arrays. Both loci have multiple pseudogenes in the Canu-only and Canu-plus Pilon assemblies, but many of these errors are removed upon supplemental correction. The presence of a pseudogene in the ME49 ROP5 locus has been predicted before based on direct sequencing, suggesting that this may represent the most accurate version of the ME49 ROP5 locus sequenced to date.

**Figure 8.**
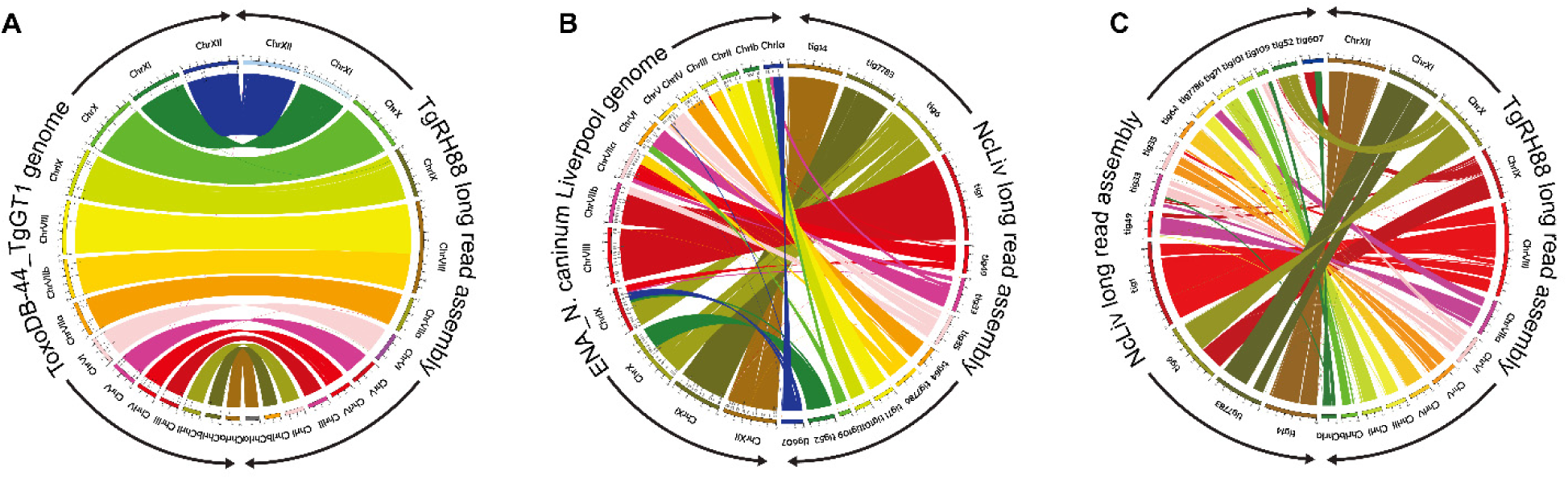
Long-read assembly reveals N. caninum karyotype and its synteny with T. gondii. (A) Circos plot showing high synteny between the TgRH88 long-read assembly and the ToxoDB-48_TgGT1 genome. (B) Circos plot showing the chromosomal translocations and inversions in NcLiv long-read assembly compared to the ENA_NcLiv genome. (C) Circos plot showing the syntenic relationship between TgRH88 and NcLiv long-read assembly.

### Long-read assembly revises *N. caninum* karyotype its lack of synteny with *T. gondii*

The comparison of *Tg*RH88 long-read assembly and annotation with the ToxoDB-48_*Tg*GT1 genome revealed a high level of collinearity between the two genomes with no large-scale rearrangement between chromosomes (except for the chrVIIb/VIII fusion) (**Figure 8A**), whereas a large number of chromosomal translocations and inversions were observed in the *Nc*Liv long-read assembly with respect to ENA_*Nc*Liv genome (**Figure 8B**). For instance, the NcLiv_tig0000052 in *Nc*Liv long-read assembly contained a portion of chrIX and a portion of chrX in the current *N. caninum* genome (**Figure 8B**). A portion of NcLiv_tig00000006 in *Nc*Liv long-read assembly was mapped to the current *N. caninum* chrX, while the remainder of NcLiv_tig00000006 was mapped to chrIX and a region of it was inverted (**Figure 8B**). In addition to this, some chromosomal regions in the *Nc*Liv long-read assembly did not show any synteny with the ENA_*Nc*Liv genome (**Figure 8B**). Similar chromosomal rearrangement patterns were observed in other *N. caninum* strains (as described in the co-submitted paper).

While previous studies (Reid *et al*. 2012; Lorenzi *et al*. 2016a) showed that the current *T. gondii* and *N. caninum* reference genomes were highly syntenic, the comparison of *Tg*RH88 long-read assembly with the *Nc*Liv long-read assembly showed smaller blocks of synteny in most regions between the two genomes (**Figure 8C**). Many individual contigs of the *Nc*Liv long-read assembly were shown to be mapped to multiple chromosome contigs in the *Tg*RH88 long-read assembly. For example, a 3.26-Mb region and a 4.43-Mb region of TgRH88_tig00000006 of the *Nc*Liv long-read assembly were in synteny with regions on TgRH88_tig00000001 and TgRH88_tig00000003 of *Tg*RH88 long-read assembly, respectively (**Figure 8C**). Moreover, some regions of *Nc*Liv long-read assembly (e.g. NcLiv_tig00007783: 2,000,000-2,500,000 bp) exhibit no synteny with the chromosomes of *Tg*RH88 genome (**Figure 8C**). Overall, our long-read assembly revealed a new and accurate *N. caninum* karyotype, and revised the genomic synteny between the two closely-related species, *T. gondii* and *N. caninum*.

## DISCUSSION

The first wave of genome sequencing using first- and second-generation sequencing technologies revolutionized our ability to link phenotype to genotype in diverse strains of *Toxoplasma gondii* and its near relatives. Sequence from multiple isolates have been made publicly available and hosted on ToxoDB.org and were outlined in a recent publication (Lorenzi *et al*. 2016a). These genomes, while of great use, were expectedly incomplete due to the presence of hundreds to thousands of sequence assembly gaps depending on sequencing depth and methodology. Recent years have experienced the impact of so-called “3rd generation” technologies that have revolutionized the speed, cost and efficacy of de novo sequence assembly, and most importantly provide a means to dramatically improve existing sequence assemblies. The critical feature of these approaches is the fact that they generate sequence read lengths hundreds to thousands of times longer than those generated using 1st and 2nd generation approaches, and can span repetitive sequence (both short tandem repeats and larger segmental duplications). This technology provides a unique opportunity to greatly improve the assembly of the *T. gondii* genome, particularly at repetitive loci which are known to encode diversified secreted effectors (Adomako-Ankomah *et al*. 2014; Adomako-Ankomah *et al*. 2016).

Our work revises the *T. gondii* and *N. caninum* karyotypes by identifying a previously unappreciated fusion between segments previously thought to be distinct chromosomes (VIIb and VIII). Throughout the history of the *T. gondii* genome the number of linkage groups has been a moving target and has become more precise as new mapping and sequencing technologies have become available. Initial genetic mapping experiments and high molecular weight Southern blotting identified 11 linkage groups (Sibley and Boothroyd 1992), while a denser map in later studies identified 13 linkage groups (Su *et al*. 2002). This particular map was still not fully representative of the *T. gondii* karyotype, as markers known to be on chromosome VI were included in linkage group X, chromosome XII was split into 2 linkage groups (“Unknown 1” and “Unknown 2”), and chromosome XI was missing (Su *et al*. 2002). A clearer picture emerged when this linkage map was integrated with shotgun sequence assembly data using 1^st^ generation sequencing, leading to a consensus karyotype of 14 chromosomes. In all three of these assemblies (from strains GT1, ME49 and VEG), chromosomes VIIb and VIII always assembled into separate contigs. However, in some studies significant genetic linkage between markers on former chromosomes VIIb and VIII was observed (for example see the discussion in (Khan *et al*. 2005) and figure S2 in (Khan *et al*. 2014)), but the lack of any contiguous assemblies for these two genome fragments (as found on ToxoDB.org as well as outlined in (Lorenzi *et al*. 2016a)) led to continued acceptance of the 14 chromosome model. It was only until recently where reports using chromosome-capture technologies (“Hi-C”; (Bunnik *et al*. 2019)) suggested a fusion between the VIIb and VIII genome fragments, and this was the first study to propose a 13 chromosome karyotype that is most consistent with our nuclear genome assembly..

Our work here conclusively establishes the *T. gondii* 13 chromosome karyotype in *T. gondii* strain ME49, and show that this karyotype is conserved across all queried *T. gondii* strains and one of its near relatives, *Neospora caninum*. The reason for the consistent prediction that fragments VIIb and VIII were distinct in *T. gondii* was clearly due to repetitive sequences near the breakpoint (hence the consistent and artificial fragmentation of this chromosome across multiple de novo sequenced strains using both first and second generation sequencing technology (Lorenzi *et al*. 2016a)). For *N. caninum*, it appears that the 14 chromosome model was based on overestimation of the degree of synteny between *T. gondii* and *N. caninum* (Reid *et al*. 2012). This karyotype that is robustly supported by our assemblies are consistent with existing Hi-C data (Bunnik *et al*. 2019) and existing genetic linkage maps from F1 progeny derived from type IxII and IxIII crosses (Khan *et al*. 2005; Khan *et al*. 2014).

Tandem gene expansion followed by selection-driven diversification provides a means for genome innovation and neofunctionalization, and this has occurred at multiple loci in the *T. gondii* genome (Adomako-Ankomah *et al*. 2014; Adomako-Ankomah *et al*. 2016; Blank and Boyle 2018). These loci can differ in copy number between strains, including those belonging to the same clonal lineage (for example MAF1 and ROP5 copy number differs between “Type 1” strains GT1 and RH (Adomako-Ankomah *et al*. 2014; Adomako-Ankomah *et al*. 2016), and MAF1 copy number differs between “Type 3” strains CTG and VEG (Adomako-Ankomah *et al*. 2016). While it is generally assumed that copy number changes can occur during errors in DNA replication, this could occur with different frequency during sexual versus asexual propagation. Here we show that some loci can change in gene number and content during sexual recombination by sequencing multiple F1 progeny from a well-defined cross between type 2 and type 3 *T. gondii*. Specific changes in copy number and/or content at specific loci could have a dramatic impact on the overall virulence phenotypes of individual F1 progeny that emerge from natural crosses.

Our long-standing interest in tandem gene arrays was one of the original motivations of this study as these loci are particularly prone to incomplete and inaccurate assembly in a variety of species including *T. gondii* and N. caninum (Adomako-Ankomah *et al*. 2014; Jain *et al*. 2018; Lapp *et al*. 2018). While the above analysis was sufficient to accurately determine the number, order and orientation of these genes in multiple strains of *T. gondii* including the F1 progeny clones, the loci were still artifactually pseudogenized even after multiple rounds of polishing with Pilon. We are not aware a comprehensive attempt to solve this problem, possibly because single copy genes tend to be very accurately corrected using tools like Pilon (Senol Cali *et al*. 2019). As a case in point, a hybrid PacBioRS/Hi-C assembly of the *P. knowlesi* genome (Lapp *et al*. 2018) required manual assembly correction to removed hundreds of incorrectly pseudogenized members of the *SICAvar* gene family. Our approach here relies on a similar approach but is automated to permit multiple rounds of error correction efficiently using a multi-core computer running Linux. Based on our analyses of these sequences they appear to be the most precise version of these intractable sequences to date, given that current versions of these genes deposited to Genbank from prior publications from our group and others on ROP5 (Reese et al. 2011) and MAF1 (Adomako-Ankomah et al. 2014; Adomako-Ankomah et al. 2016) were obtained by PCR and subject to artefactual chimerism. This is particularly evident for the MAF1 locus which is made up of tandem expansions of a 2 gene cassette (which we have dubbed MAF1A and MAF1B and these genes are functionally distinct (only MAF1B drives the unique T. gondii host mitochondrial association phenotype (Adomako-Ankomah et al. 2016; Blank et al. 2018)).

Taken together these data clearly demonstrate that the *T. gondii* karyotype has been mischaracterized as being made up of 14, rather than 13, chromosomes. The 13 chromosome karyotype is consistent across multiple strains and is also conserved across the local phylogeny, in that *N. caninum* has the same number. This is consistent with a propensity for chromosome number to be largely conserved between closely-related species even when gene content and overall synteny may differ dramatically (as they do for *N. caninum* and *T. gondii*, in contrast to previous work that overestimated the extent of synteny between these species; (Reid *et al*. 2012)). These data represent a new era in genome sequencing in *T. gondii* and its near relatives, allowing for near-complete telomere-to-telomere assemblies of *T. gondii* strains to be generated with minimal effort and cost. Moreover with existing and supplementary correction methods that are targeted to the systematic error that is most common in Oxford Nanopore sequencing data (incorrect calls of homopolymer nucleotide runs), we have been able to generate the most accurate version of a key subset of virulence effector genes in this organism with such an enormous impact on human global heatlh.

## METHODS

### Parasite and cell culture

All *T. gondii* strains and *N. caninum* Liverpool strain were maintained by serial passage of tachyzoites in human foreskin fibroblasts (HFFs). HFFs were cultured in Dulbecco’s modified Eagle’s medium (DMEM) containing 10% fetal bovine serum (FBS), 2 mM glutamine, and 50 mg/ml each of penicillin and streptomycin at 37 °C in a 5% CO_2_ incubator.

### High-molecular-weight (HMW) genomic DNA extraction

Prior to DNA purification, tachyzoites of *T. gondii* or *N. caninum* Liverpool strain were grown in 2×10^7^ HFFs for about 5-7 days until the monolayer was fully infected. The infected cells were then scraped, and syringe-lysed to release the parasites, and the parasites were harvested by filtering (5.0 μm syringe filter, Millipore) and centrifugation. The pelleted parasites were resuspended and lysed in 10 ml TLB buffer (100 mM NaCl, 10 mM Tris-HCl [pH 8.0], 25 mM EDTA [pH 8.0], 0.5% (w/v) SDS) containing 20 μg/ml RNase A for 1 hour at 37 °C, followed by a 3-hour proteinase K (20 mg/ml) digestion at 50 °C. The lysate was split into two tubes containing phase-lock gel (Quantabio), and 5 ml TE-saturated phenol (Millipore Sigma) was added to each tube, mixed by rotation for 10 min and centrifuged for 10 min at 4,750×g. The DNA was isolated by removing the aqueous phase to two tubes containing phase-lock gel, followed by a 25:24:1 phenol-chloroform-isoamyl alcohol (Millipore Sigma) extraction. The DNA in the aqueous phase was further purified by ethanol precipitation by adding 4 ml 3 M NaOAc (pH 5.2), and then mixing 30 ml ice-cold 100% ethanol. The solution was mixed by gentle inversion and briefly centrifuged at 1000×g for 2 min to pellet the DNA. The resulting pellet was washed three times with 70% ethanol, and all visible traces of ethanol were removed from the tube. The DNA was allowed to air dry for 5 min on a 40 °C heat block, resuspended in 40 μl elution buffer (10 mM Tris-HCl [pH 8.5]) without mixing on pipetting, followed by an overnight incubation at 4 °C. The concentration and purity of the eluted DNA were measured using a NanoDrop spectrophotometer (Thermo Scientific), and approximately 400 ng of DNA was used for sequencing library preparation.

### MinION library preparation and sequencing

The MinION sequencing libraries were prepared using the SQK-RAD004 or SQK-RBK004 kit (Oxford Nanopore Technologies) protocol accompanying all pipetting steps performed using pipette tips with ∼1 cm cut off of the end. High molecular weight DNA (7.5 μl corresponding to 400ng of DNA) was mixed with 2.5 μl of fragmentation mix (SQK-RAD004 kit) or barcoded fragmentation mix (SQK-RBK004 kit), and then incubated at 30 °C for 1 min, followed by 80 °C for 1 min on a thermocycler. After incubation, 1 μl of rapid adapter mix was added and mixed gently by flicking the tube, and the library was incubated at room temperature for 5 min. Prior to the library loading, the flow cell (MinION R9.4.1 flow cell [FLO-MIN106, Oxford Nanopore Technologies]) was primed by loading 800 μl of priming mix (flush tether and flush buffer mix, Oxford Nanopore Technologies) into the priming port on the flow cell and left for 5 min. After priming, 11 μl of DNA library was mixed with 34 μl of sequencing buffer (Oxford Nanopore Technologies), 25.5 μl of resuspended loading beads (Oxford Nanopore Technologies), and 4.5 μl of nuclease-free water. To initiate sequencing, 75 μl of the prepared library was loaded onto the flow cell through the SpotON sample port in a drop-by-drop manner. Sequencing was performed immediately after platform QC, which determined the number of active pores. The sequencing process was controlled using MinKNOW (Oxford Nanopore Technologies) and the resulting FAST5 files were base-called using Guppy v3.2.1 (Oxford Nanopore Technologies). The barcoded sequencing reads were demultiplexed using Deepbinner (https://github.com/rrwick/Deepbinner). Read statistics were computed and graphed using Nanoplot v1.0.0 (De Coster *et al*. 2018).

### Read quality control and *de novo* genome assembly

To assess read quality, raw sequencing reads were aligned against the reference genomes (Information of the reference genomes used in this study are shown in **Table S1**) using Minimap2 (Li 2018) with the following parameter: -ax map-ont. All reads > 1000 bp in length were input into Canu v1.7.1 (Koren *et al*. 2017) for *de novo* assembly using the complete Canu pipeline (correction, trimming, and assembly) (Koren *et al*. 2017) with the following parameters: correctedErrorRate=0.154, gnuplotTested=TRUE, minReadLength=1000, and –nanopore-raw. Assembly was performed based on an estimated 65 Mb genome size for *T. gondii* strains, and 57 Mb for *N. caninum* Liverpool strain, and was run using the Slurm management system on the high throughput computing (HTC) cluster at University of Pittsburgh.

### Error correction and assembly polishing

For the Canu-yielded *Tg*RH88, *Tg*ME49, *Tg*CTG, and *Nc*Liv assemblies, assembly errors were corrected by Pilon v1.23 (Walker *et al*. 2014) with four iterations using the alignment of select whole-genome Illumina paired-end reads (SRA: SRR5123638, SRR2068653, SRR5643140, or ERR701181) to the assembly contigs generated by BWA-MEM (Li 2013). The resulting corrected contigs were reassembled using Flye v2.5 (Kolmogorov *et al*. 2019). For the II×III F1 progeny assemblies, CL13, S27, S21, S26, and D3X1, the assembly contigs were directly subjected to Flye for reassembly without Pilon correction. The final contigs/scaffolds in the *Tg*RH88, *Tg*ME49, and *Tg*CTG assemblies were assigned, ordered, and oriented to chromosomes using ToxoDB-48 genomes as reference (**Table S1**). These genomes were then deposited in Genbank.

### Supplemental assembly correction

Since tandem gene arrays were still artificially pseudogenized after Pilon-based error correction we performed multiple rounds of error correction to eliminate any remaining homopolymer errors. First we aligned the Illumina sequence reads used in Pilon to the polished assembly using bowtie2 (using the -k 10 parameter to allow reads to map to up to 10 distinct locations). We then broke the genome into 250 Kb fragments and counted all possible 30 bp kmers in the raw reads aligning to that 250 Kb region. Then, we walked through the assembly 1 bp at a time, extracting the 30 bp starting at that position and counting the number of times that 30 bp was found in the reads mapping to the 250 Kb region. If the read count was < 5 we attempted to determine the length of the homopolymer by adding sequence iteratively until the number of reads harboring that sequence increased above 10. The nucleotide to be added was selected only of ≥90% of the reads had the same nucleotide at that position. If the read was not improved after addition of 10 nucleotides, no changes were made to the sequence, unless the truncation of 1 nucleotide from the end of the 30 bp assembly fragment increased coverage, in which case that change was made to the assembly. We repeated this correction 4 times iteratively, and used the corrected assemblies to determine the sequence of individual paralogs at tandem gene arrays such as MAF1, ROP5 and ROP38.

### Long-read assembly evaluation

Assembly statistics were computed using Canu v1.7.1 and QUAST v5.0.2 (Mikheenko *et al*. 2018). Genome assembly completeness assessment was performed using BUSCO v3.0.2 (Waterhouse *et al*.) against the Protists ensembl dataset. Gene predictions were performed using Augustus v3.3 (Keller *et al*. 2011) with the *Toxoplasma gondii*-specific training set.

### Whole genome alignment

Whole genome alignments between the long-read assemblies and the reference genomes were performed using MUMmer v4.0.0 (Kurtz *et al*. 2004) and Mauve v2.4.0 (Darling *et al*. 2004). Dotplots were generated using D-Genies (Cabanettes and Klopp 2018). BWA-MEM was used for remapping the corrected reads to the reference genomes, and all SAM files were parsed to sorted BAM files using SAMtools v1.9 (Li *et al*. 2009). Alignments were visualized using a variety of tools including IGV v2.4.15 (Thorvaldsdottir *et al*. 2013), Mauve (Darling *et al*. 2004), Circos (Krzywinski *et al*. 2009) and custom scripts implemented in R statistical software (Team 2020).

### Structural variant detection

Structural variant differences between the long-read assemblies and reference genomes (from ToxoDB.org or Genbank, depending on the strains analyzed) were identified by processing the delta file generated by the MUMmer alignment generator NUCmer with the parameter “show-diff”. In addition, manual curation of structural variants was performed by visual inspection of chromosomal rearrangements based on the whole genome alignments generated using Mauve and MUMmer, and using BLASTN to identify and count repetitive loci. Select alignment plots were generated to integrate these data using R statistical software (Team 2020).

### Copy number variant detection

For all gene-coding tandem expansions in *T. gondii* or *N. caninum* identified previously (Adomako-Ankomah *et al*. 2014), all predicted gene sequences were downloaded from ToxoDB.org and aligned using BLASTN, and alignments showing > 80% identity and covering at least 80% of the query gene were used to count tandem repeat numbers in *T. gondii* strains RH88, ME49 and CTG. For a subset of these tandemly expanded loci (*ROP5*, *ROP38*, *MIC17*, *MAF1*, *ROP4/7*, and *TSEL8*), similar analyses were performed manually against all queried *T. gondii* strains (including F1 progeny clones), and in this case only alignments that showed more than 95% identity and 98% coverage were considered as a match. Paralogs were grouped based on alignment identity, and the number of copies at these loci was estimated by alignment match counts. Only matches that were within a single assembled Nanopore-derived contig were considered for copy number estimates, and the length of the sequence between the upstream of the first match or the downstream of the last match on the genomic coordinate and the edge of the corresponding contig had to be longer than that of the sequence between two adjacent matches.

### Identification and analysis of new sequences in the long-read assemblies

To identify new sequences that filled reference assembly gaps, we aligned the long-read chromosome contigs to the reference assembly chromosomes using NUCmer with the “show-diff -q” parameter. The coordinates of a) sequence expansions and b) unaligned sequences from our *de novo* assembly were determined, and the corresponding sequences were extracted using custom scripts. Repeats in all genome assemblies were detected using Tandem Repeat Finder (TRF) v4.09 (Benson 1999) and all repeats with a period size of ≥ 500 bp and having at least 2 copies were used to determine the impact of long read assembly on resolution of repeats > 500 bp in size (which are poorly resolved by 2^nd^ or 3^rd^ generation sequencing technologies.

### Hi-C Data analysis

Published Hi-C reads (Bunnik *et al*. 2019) were realigned to assemblies TgRH88, TgME49, TgCTG, S27, and S21, then processed further by assigning fragments and removing invalid and duplicate pairs using the processing pipeline HiCPro (Servant *et al*. 2015). Resulting raw intrachromosomal and interchromosomal contact maps were built at 10-kb resolution and corrected for experimental and technical biases using ICE normalization (Imakaev *et al*. 2012).

## DATA ACCESS

All raw sequence reads (as fastq files) and polished assemblies have been deposited at Genbank under BioProject number PRJNA638608.

## ACKNOWLEDGEMENTS

The authors would like to thank members of the Boyle lab for critical reading of the manuscript and Josh Quick for public sharing of protocols for DNA isolation that maximize read length. This work was supported by grants R01AI116855 and R01AI114655 to J.P.B., State 554 Scholarship Fund from the China Scholar Council (201708440340) to JX, grant R21 AI142506 (NIH) and NIFA-Hatch-225935 (University of California, Riverside) to KGLR.

## REFERENCES

Adomako-Ankomah Y, English ED, Danielson JJ, Pernas LF, Parker ML, Boulanger MJ, Dubey JP, Boyle JP. 2016. Host Mitochondrial Association Evolved in the Human Parasite Toxoplasma gondii via Neofunctionalization of a Gene Duplicate. Genetics 203: 283–298.

Adomako-Ankomah Y, Wier GM, Borges AL, Wand HE, Boyle JP. 2014. Differential locus expansion distinguishes Toxoplasmatinae species and closely related strains of Toxoplasma gondii. mBio 5: e01003–01013.

Benson G. 1999. Tandem repeats finder: a program to analyze DNA sequences. Nucleic acids research 27: 573–580.

Blank ML, Boyle JP. 2018. Effector variation at tandem gene arrays in tissue-dwelling coccidia: who needs antigenic variation anyway? Current opinion in microbiology 46: 86–92.

Blank ML, Parker ML, Ramaswamy R, Powell CJ, English ED, Adomako-Ankomah Y, Pernas LF, Workman SD, Boothroyd JC, Boulanger MJ et al. 2018. A Toxoplasma gondii locus required for the direct manipulation of host mitochondria has maintained multiple ancestral functions. Mol Microbiol 108: 519–535.

Brooks CF, Francia ME, Gissot M, Croken MM, Kim K, Striepen B. 2011. Toxoplasma gondii sequesters centromeres to a specific nuclear region throughout the cell cycle. Proceedings of the National Academy of Sciences of the United States of America 108: 3767–3772.

Bunnik EM, Venkat A, Shao J, McGovern KE, Batugedara G, Worth D, Prudhomme J, Lapp SA, Andolina C, Ross LS et al. 2019. Comparative 3D genome organization in apicomplexan parasites. Proceedings of the National Academy of Sciences of the United States of America doi:10.1073/pnas.1810815116.

Burg JL, Grover CM, Pouletty P, Boothroyd JC. 1989. Direct and sensitive detection of a pathogenic protozoan, Toxoplasma gondii, by polymerase chain reaction. Journal of clinical microbiology 27: 1787–1792.

Cabanettes F, Klopp C. 2018. D-GENIES: dot plot large genomes in an interactive, efficient and simple way. PeerJ 6: e4958.

Clemente M, de Miguel N, Lia VV, Matrajt M, Angel SO. 2004. Structure analysis of two Toxoplasma gondii and Neospora caninum satellite DNA families and evolution of their common monomeric sequence. J Mol Evol 58: 557–567.

Darling AC, Mau B, Blattner FR, Perna NT. 2004. Mauve: multiple alignment of conserved genomic sequence with rearrangements. Genome Res 14: 1394–1403.

De Coster W, D’Hert S, Schultz DT, Cruts M, Van Broeckhoven C. 2018. NanoPack: visualizing and processing long-read sequencing data. Bioinformatics (Oxford, England) 34: 2666–2669.

Delcher AL, Salzberg SL, Phillippy AM. 2003. Using MUMmer to identify similar regions in large sequence sets. Curr Protoc Bioinformatics Chapter 10: Unit 10 13.

Diaz-Viraque F, Pita S, Greif G, Moreira de Souza RdC, Iraola G, Robello C. 2018. Nanopore sequencing significantly improves genome assembly of the eukaryotic protozoan parasite Trypanosoma cruzi. bioRxiv doi:10.1101/489534: 489534.

Dudchenko O, Batra SS, Omer AD, Nyquist SK, Hoeger M, Durand NC, Shamim MS, Machol I, Lander ES, Aiden AP et al. 2017. De novo assembly of the Aedes aegypti genome using Hi-C yields chromosome-length scaffolds. Science 356: 92–95.

Echeverria PC, Rojas PA, Martin V, Guarnera EA, Pszenny V, Angel SO. 2000. Characterisation of a novel interspersed Toxoplasma gondii DNA repeat with potential uses for PCR diagnosis and PCR-RFLP analysis. FEMS Microbiol Lett 184: 23–27.

Edvinsson B, Lappalainen M, Evengard B, Toxoplasmosis ESGf. 2006. Real-time PCR targeting a 529-bp repeat element for diagnosis of toxoplasmosis. Clinical microbiology and infection : the official publication of the European Society of Clinical Microbiology and Infectious Diseases 12: 131–136.

Ewing B, Green P. 1998. Base-calling of automated sequencer traces using phred. II. Error probabilities. Genome Res 8: 186–194.

Ewing B, Hillier L, Wendl MC, Green P. 1998. Base-calling of automated sequencer traces using phred. I. Accuracy assessment. Genome Res 8: 175–185.

Fournier T, Gounot JS, Freel K, Cruaud C, Lemainque A, Aury JM, Wincker P, Schacherer J, Friedrich A. 2017. High-Quality de Novo Genome Assembly of the Dekkera bruxellensis Yeast Using Nanopore MinION Sequencing. G3 (Bethesda, Md) 7: 3243–3250.

Gajria B, Bahl A, Brestelli J, Dommer J, Fischer S, Gao X, Heiges M, Iodice J, Kissinger JC, Mackey AJ et al. 2008. ToxoDB: an integrated Toxoplasma gondii database resource. Nucleic acids research 36: D553–556.

Gissot M, Walker R, Delhaye S, Huot L, Hot D, Tomavo S. 2012. Toxoplasma gondii chromodomain protein 1 binds to heterochromatin and colocalises with centromeres and telomeres at the nuclear periphery. PloS one 7: e32671.

Imakaev M, Fudenberg G, McCord RP, Naumova N, Goloborodko A, Lajoie BR, Dekker J, Mirny LA. 2012. Iterative correction of Hi-C data reveals hallmarks of chromosome organization. Nature methods 9: 999–1003.

Jain M, Koren S, Miga KH, Quick J, Rand AC, Sasani TA, Tyson JR, Beggs AD, Dilthey AT, Fiddes IT et al. 2018. Nanopore sequencing and assembly of a human genome with ultra-long reads. Nature biotechnology 36: 338–345.

Jones JL, Holland GN. 2010. Annual burden of ocular toxoplasmosis in the US. The American journal of tropical medicine and hygiene 82: 464–465.

Joynson DH, Wreghitt TG. 2005. Toxoplasmosis: a comprehensive clinical guide. Cambridge University Press.

Keller O, Kollmar M, Stanke M, Waack S. 2011. A novel hybrid gene prediction method employing protein multiple sequence alignments. Bioinformatics (Oxford, England) 27: 757–763.

Khan A, Behnke MS, Dunay IR, White MW, Sibley LD. 2009. Phenotypic and gene expression changes among clonal type I strains of Toxoplasma gondii. Eukaryotic cell 8: 1828–1836.

Khan A, Shaik JS, Behnke M, Wang Q, Dubey JP, Lorenzi HA, Ajioka JW, Rosenthal BM, Sibley LD. 2014. NextGen sequencing reveals short double crossovers contribute disproportionately to genetic diversity in Toxoplasma gondii. BMC Genomics 15: 1168.

Khan A, Taylor S, Su C, Mackey AJ, Boyle J, Cole R, Glover D, Tang K, Paulsen IT, Berriman M et al. 2005. Composite genome map and recombination parameters derived from three archetypal lineages of Toxoplasma gondii. Nucleic acids research 33: 2980–2992.

Kim K, Weiss LM. 2004. Toxoplasma gondii: the model apicomplexan. International journal for parasitology 34: 423–432.

Kolmogorov M, Yuan J, Lin Y, Pevzner PA. 2019. Assembly of long, error-prone reads using repeat graphs. Nature biotechnology 37: 540–546.

Koren S, Walenz BP, Berlin K, Miller JR, Bergman NH, Phillippy AM. 2017. Canu: scalable and accurate long-read assembly via adaptive k-mer weighting and repeat separation. Genome Res 27: 722–736.

Krzywinski M, Schein J, Birol I, Connors J, Gascoyne R, Horsman D, Jones SJ, Marra MA. 2009. Circos: an information aesthetic for comparative genomics. Genome Res 19: 1639–1645.

Kurtz S, Phillippy A, Delcher AL, Smoot M, Shumway M, Antonescu C, Salzberg SL. 2004. Versatile and open software for comparing large genomes. Genome biology 5: R12.

Lapp SA, Geraldo JA, Chien JT, Ay F, Pakala SB, Batugedara G, Humphrey J, Ma Hc, De BJ, Le Roch KG et al. 2018. PacBio assembly of a Plasmodium knowlesi genome sequence with Hi-C correction and manual annotation of the SICAvar gene family. Parasitology 145: 71–84.

Lau YL, Lee WC, Gudimella R, Zhang G, Ching XT, Razali R, Aziz F, Anwar A, Fong MY. 2016. Deciphering the Draft Genome of Toxoplasma gondii RH Strain. PloS one 11: e0157901.

Li H. 2013. Aligning sequence reads, clone sequences and assembly contigs with BWA-MEM. arXiv:13033997v2 [q-bioGN].

Li H. 2018. Minimap2: pairwise alignment for nucleotide sequences. Bioinformatics (Oxford, England) 34: 3094–3100.

Li H, Handsaker B, Wysoker A, Fennell T, Ruan J, Homer N, Marth G, Abecasis G, Durbin R. 2009. The Sequence Alignment/Map format and SAMtools. Bioinformatics (Oxford, England) 25: 2078–2079.

Lorenzi H, Khan A, Behnke MS, Namasivayam S, Swapna LS, Hadjithomas M, Karamycheva S, Pinney D, Brunk BP, Ajioka JW et al. 2016a. Local admixture of amplified and diversified secreted pathogenesis determinants shapes mosaic Toxoplasma gondii genomes. Nature Communications 7: 10147.

Lorenzi H, Khan A, Behnke MS, Namasivayam S, Swapna LS, Hadjithomas M, Karamycheva S, Pinney D, Brunk BP, Ajioka JW et al. 2016b. Local admixture of amplified and diversified secreted pathogenesis determinants shapes mosaic Toxoplasma gondii genomes. Nature Communications 7: 10147.

Madoui MA, Engelen S, Cruaud C, Belser C, Bertrand L, Alberti A, Lemainque A, Wincker P, Aury JM. 2015. Genome assembly using Nanopore-guided long and error-free DNA reads. BMC Genomics 16: 327.

Matrajt M, Angel SO, Pszenny V, Guarnera E, Roos DS, Garberi JC. 1999. Arrays of repetitive DNA elements in the largest chromosomes of Toxoplasma gondii. Genome 42: 265–269.

Michael TP, Jupe F, Bemm F, Motley ST, Sandoval JP, Lanz C, Loudet O, Weigel D, Ecker JR. 2018. High contiguity Arabidopsis thaliana genome assembly with a single nanopore flow cell. Nat Commun 9: 541.

Mikheenko A, Prjibelski A, Saveliev V, Antipov D, Gurevich A. 2018. Versatile genome assembly evaluation with QUAST-LG. Bioinformatics (Oxford, England) 34: i142–i150.

Pfefferkorn ER, Pfefferkorn LC. 1976. Toxoplasma gondii: isolation and preliminary characterization of temperature-sensitive mutants. Experimental parasitology 39: 365–376.

Quick J. 2018. Ultra-long read sequencing protocol for RAD004 V.3. https://www.protocols.io/view/ultra-long-read-sequencing-protocol-for-rad004-mrxc57n?version_warning=no.

Reese ML, Zeiner GM, Saeij JP, Boothroyd JC, Boyle JP. 2011. Polymorphic family of injected pseudokinases is paramount in Toxoplasma virulence. Proceedings of the National Academy of Sciences of the United States of America 108: 9625–9630.

Reid AJ, Vermont SJ, Cotton JA, Harris D, Hill-Cawthorne GA, Könen-Waisman S, Latham SM, Mourier T, Norton R, Quail MA et al. 2012. Comparative Genomics of the Apicomplexan Parasites Toxoplasma gondii and Neospora caninum: Coccidia Differing in Host Range and Transmission Strategy. PLOS Pathogens 8: e1002567.

Reischl U, Bretagne S, Kruger D, Ernault P, Costa JM. 2003. Comparison of two DNA targets for the diagnosis of Toxoplasmosis by real-time PCR using fluorescence resonance energy transfer hybridization probes. BMC infectious diseases 3: 7.

Saeij JP, Boyle JP, Boothroyd JC. 2005. Differences among the three major strains of Toxoplasma gondii and their specific interactions with the infected host. Trends Parasitol 21: 476–481.

Sanger F, Nicklen S, Coulson AR. 1977. DNA sequencing with chain-terminating inhibitors. Proceedings of the National Academy of Sciences of the United States of America 74: 5463–5467.

Schmidt MH, Vogel A, Denton AK, Istace B, Wormit A, van de Geest H, Bolger ME, Alseekh S, Mass J, Pfaff C et al. 2017. De Novo Assembly of a New Solanum pennellii Accession Using Nanopore Sequencing. The Plant cell 29: 2336–2348.

Senol Cali D, Kim JS, Ghose S, Alkan C, Mutlu O. 2019. Nanopore sequencing technology and tools for genome assembly: computational analysis of the current state, bottlenecks and future directions. Briefings in bioinformatics 20: 1542–1559.

Servant N, Varoquaux N, Lajoie BR, Viara E, Chen CJ, Vert JP, Heard E, Dekker J, Barillot E. 2015. HiC-Pro: an optimized and flexible pipeline for Hi-C data processing. Genome biology 16: 259.

Sibley LD, Ajioka JW. 2008. Population structure of Toxoplasma gondii: clonal expansion driven by infrequent recombination and selective sweeps. Annual review of microbiology 62: 329–351.

Sibley LD, Boothroyd JC. 1992. Construction of a molecular karyotype for Toxoplasma gondii. Molecular and biochemical parasitology 51: 291–300.

Su C, Howe DK, Dubey JP, Ajioka JW, Sibley LD. 2002. Identification of quantitative trait loci controlling acute virulence in Toxoplasma gondii. Proceedings of the National Academy of Sciences of the United States of America 99: 10753–10758.

Team RC. 2020. R: A language and environment for statistical computing. R Foundation for Statistical Computing, Vienna, Austria.

Thorvaldsdottir H, Robinson JT, Mesirov JP. 2013. Integrative Genomics Viewer (IGV): high-performance genomics data visualization and exploration. Briefings in bioinformatics 14: 178–192.

Villard O, Candolfi E, Ferguson DJ, Marcellin L, Kien T. 1997. Loss of oral infectivity of tissue cysts of Toxoplasma gondii RH strain to outbred Swiss Webster mice. International journal for parasitology 27: 1555–1559.

Vollger MR, Logsdon GA, Audano PA, Sulovari A, Porubsky D, Peluso P, Wenger AM, Concepcion GT, Kronenberg ZN, Munson KM et al. 2019. Improved assembly and variant detection of a haploid human genome using single-molecule, high-fidelity long reads. Annals of human genetics doi:10.1111/ahg.12364.

Walker BJ, Abeel T, Shea T, Priest M, Abouelliel A, Sakthikumar S, Cuomo CA, Zeng Q, Wortman J, Young SK et al. 2014. Pilon: An Integrated Tool for Comprehensive Microbial Variant Detection and Genome Assembly Improvement. PloS one 9: e112963.

Waterhouse RM, Seppey M, Simao FA, Manni M, Ioannidis P, Klioutchnikov G, Kriventseva EV, Zdobnov EM. BUSCO Applications from Quality Assessments to Gene Prediction and Phylogenomics.

