## Supplemental Figures and Tables for "Third generation sequencing revises the molecular karyotype for *Toxoplasma gondii* and identifies emerging copy number variants in sexual recombinants"

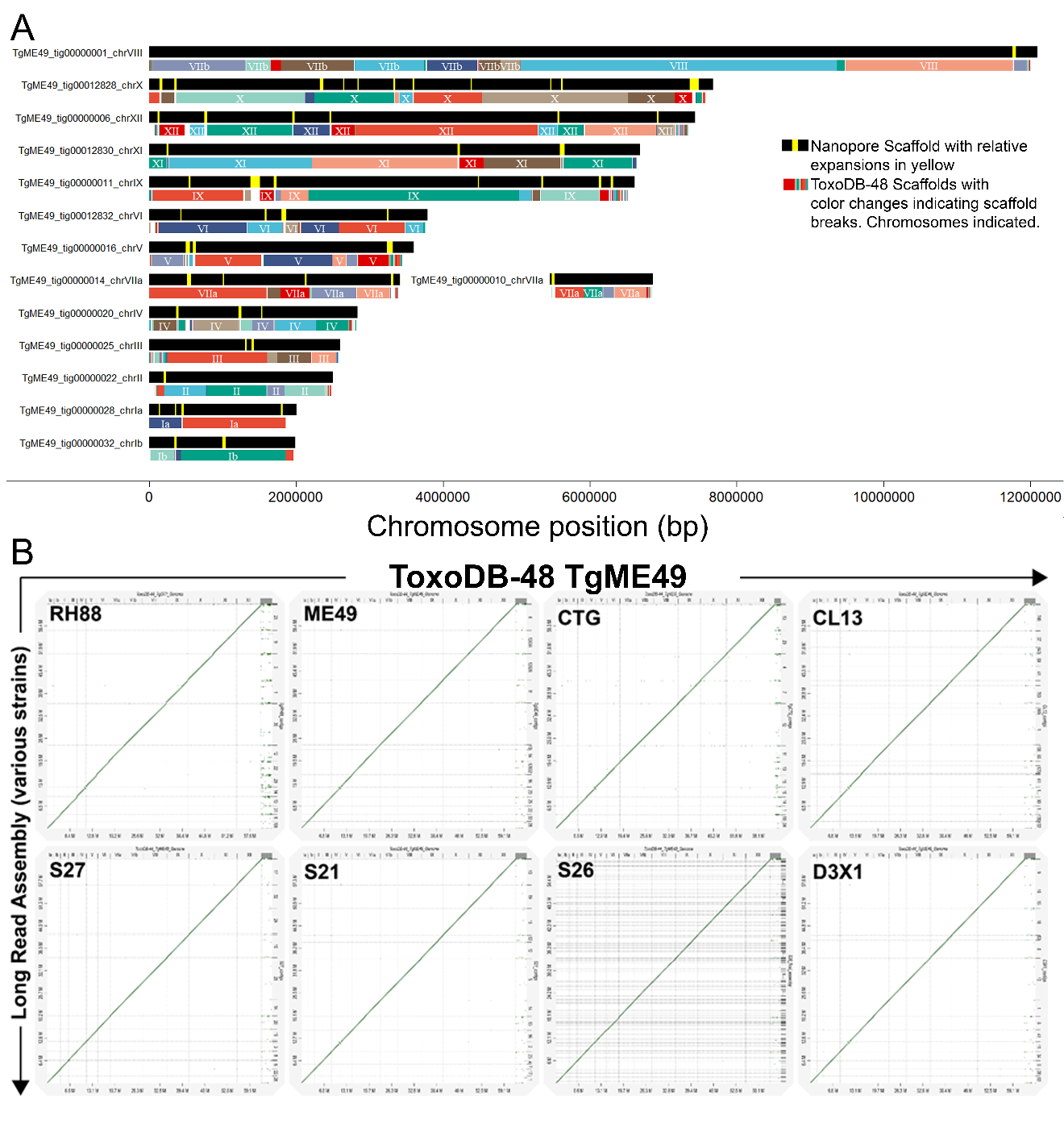

**Figure S1. A)** Positions of all contigs from the v48 release of the *T. gondii* ME49 genome mapped to the Nanopore-based *de novo* assembly. Coordinates were generated using Minimap2 and colored randomly for clarity. The chromosomal origin for each is indicated for most contigs. All chromosomes were near-complete with the exception of chromosome VIIa, which was found on 2 contigs. Yellow bars indicate genome expansions in the long-read assembly compared to the reference. **B)** Dot plot comparison of each *T. gondii* long-read assembly with the *T. gondii* ME49 genome. Plots were generated using nucmer and show high large scale synteny between all new genome assemblies and the reference.

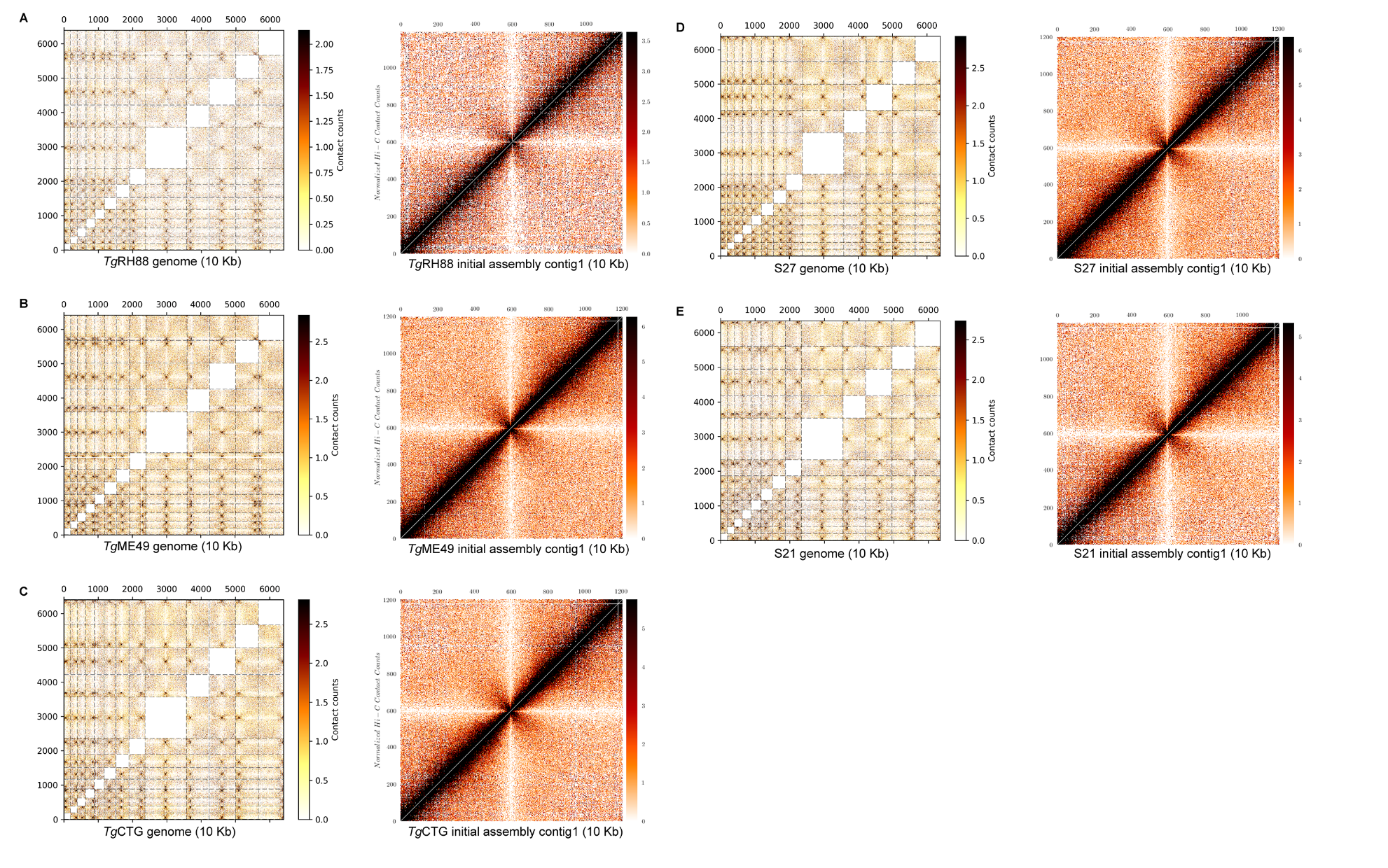
**Figure S2. Hi-C contact count heatmaps generated by aligning Hi-C reads to the long-read assembly sequences for 5 *T. gondii* genomes.** Interchromosomal contact maps are shown on the left, while the intrachromsomal contact map for the chromosome VIIb/VIII fusion (“contig 1” in all assemblies which are all named Tg[strain name]_tig00000001 in the final assemblies submitted to Genbank) is shown on the right.

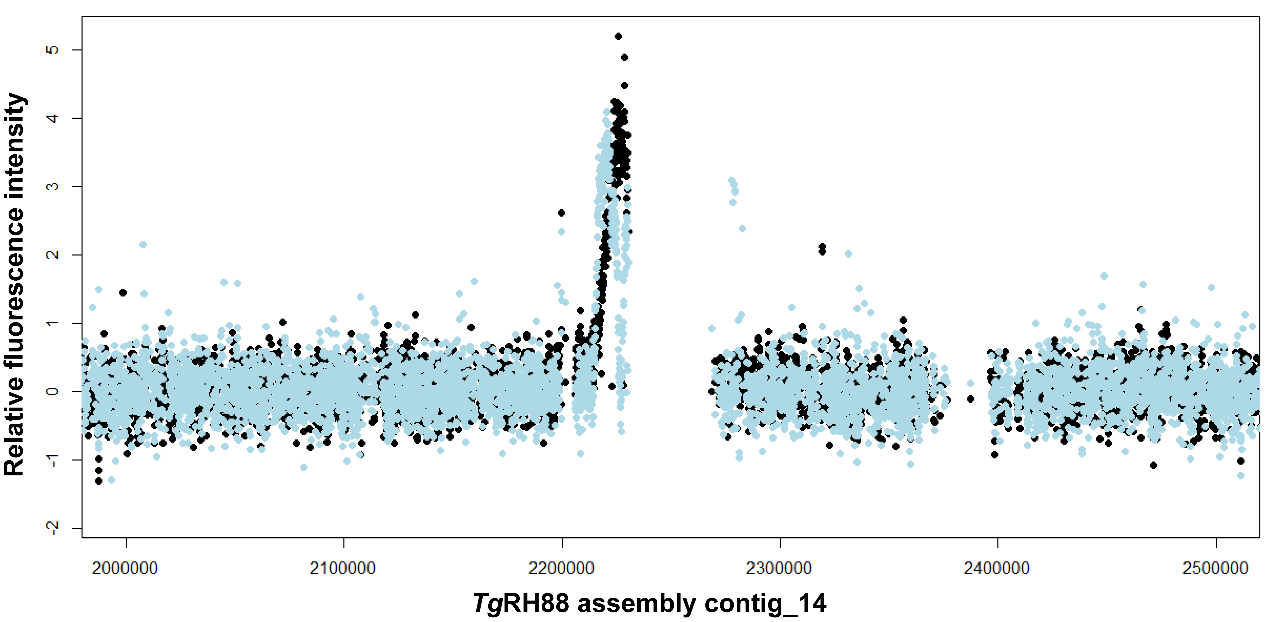

**Figure S3. Centromere identification in chromosome IV of the *T. gondii* RH88 Nanopore-based assembly.** Plot is a zoomed-in view of the ChIP-on-chip signal of centromeric histone 3 variant (CenH3) taken from (Brooks *et al.* 2011) and that shown in **Figure 4C**.

**Table S1. Description of the reference genomes used in this study.**

| **Species** | **Long-read assembly** | **Genotype (ToxoDB PCR-RFLP genotype)** | **References genome** |
| --- | --- | --- | --- |
| *Toxoplasma gondii* | *Tg*RH88 | Type I (ToxoDB #10, Type I) | ToxoDB-44_T*gondii*GT1_Genome |
| *Toxoplasma gondii* | *Tg*ME49 | Type II (ToxoDB #1, Type II) | ToxoDB-44_T*gondii*ME49_Genome |
| *Toxoplasma gondii* | *Tg*CTG | Type III (ToxoDB #2, Type III) | ToxoDB-44_T*gondii*VEG_Genome |
| *Toxoplasma gondii* | CL13 | Types II×III F1 progeny | ToxoDB-44_T*gondii*ME49_Genome or/and ToxoDB-44_T*gondii*VEG_Genome |
| *Toxoplasma gondii* | S27 | Types II×III F1 progeny |  |
| *Toxoplasma gondii* | S21 | Types II×III F1 progeny |  |
| *Toxoplasma gondii* | S26 | Types II×III F1 progeny |  |
| *Toxoplasma gondii* | D3X1 | Types II×III F1 progeny |  |
| *Neospora caninum* | *Nc*Liv | - | ENA_NcLiv |

**Table S2. Summary of sequencing statistics.**

|  | ***Tg*RH88** | ***Tg*ME49** | ***Tg*CTG** | **F1_CL13** | **F1_S27** | **F1_S21** | **F1_S26** | **F1_D3X1** | ***Neospora caninum* Liverpool** |
| --- | --- | --- | --- | --- | --- | --- | --- | --- | --- |
| # reads | 648,491 | 119,855 | 660,159 | 304,917 | 165,999 | 75,088 | 48,339 | 94,859 | 335,193 |
| Total bases (Kbp) | 7,403,277 | 2,062,812 | 7,440,130 | 3,572,013 | 2,147,437 | 1,277,107 | 607,739 | 1,319,594 | 3,368,213 |
| Coverage | 113.9× | 31.7× | 114.5× | 54.9× | 33.0× | 19.6× | 9.3× | 20.3× | 59.1× |
| Maximum read length (bp) | 158,740 | 229,201 | 175,612 | 183,371 | 186,333 | 266,564 | 152,085 | 184,336 | 116,661 |
| Read length N50 (bp) | 20,827 | 39,666 | 21,333 | 20,630 | 23,403 | 31,599 | 22,267 | 25,698 | 18,368 |
| Mean Phred score | 8.9 | 10.4 | 9.6 | 10.8 | 10.7 | 10.5 | 10.4 | 10.4 | 10.6 |
| Mapped reads (%) | 99.55 | 91.35 | 94.31 | 98.95 | 97.73 | 98.60 | 97.27 | 93.56 | 85.15 |
| Mean percent identity (%) | 83.7 | 86.2 | 86.3 | 89.0 | 88.0 | 87.3 | 86.6 | 87.0 | 87.3 |

**Table S3. Summary of read statistics after curation and correction using Canu.**

|  | ***Tg*RH88** | ***Tg*ME49** | ***Tg*CTG** | **F1_CL13** | **F1_S27** | **F1_S21** | **F1_S26** | **F1_D3X1** | ***Neospora caninum* Liverpool** |
| --- | --- | --- | --- | --- | --- | --- | --- | --- | --- |
| # corrected reads | 69,469 | 91,193 | 63,310 | 115,700 | 114,451 | 58,035 | 40,011 | 109,846 | 90,972 |
| Total corrected read bases (Kbp) | 2,637,598 | 2,637,233 | 2,609,200 | 2,610,987 | 1,687,112 | 1,076,453 | 546,526 | 1,680,973 | 2,291,712 |
| Coverage | 40.58× | 40.57× | 40.14× | 40.17× | 25.96× | 16.56× | 8.41× | 25.86× | 40.21× |
| Corrected read length N50 (bp) | 38,015 | 39,894 | 40,862 | 25,346 | 23,877 | 31,838 | 22,138 | 24,818 | 26,292 |
| Mapped reads (%) | 99.97 | 99.69 | 99.49 | 99.95 | 99.79 | 99.25 | 99.48 | 99.79 | 98.07 |
| Mean percent identity (%) | 92.9 | 95.6 | 93.4 | 97.0 | 96.2 | 94.5 | 92.7 | 96.1 | 95.1 |

**Table S4. Metrics of the initial genome assemblies yielded by Canu.**

|  | ***Tg*RH88** | ***Tg*ME49** | ***Tg*CTG** | **F1_CL13** | **F1_S27** | **F1_S21** | **F1_S26** | **F1_D3X1** | ***Neospora caninum* Liverpool** |
| --- | --- | --- | --- | --- | --- | --- | --- | --- | --- |
| # contigs | 23 | 38 | 38 | 36 | 69 | 39 | 239 | 49 | 58 |
| Total bases (bp) | 64,401,064 | 64,522,756 | 64,789,158 | 64,701,501 | 64,573,169 | 63,770,490 | 60,576,027 | 64,362,372 | 61,799,150 |
| Maximum contig length (bp) | 11,930,269 | 12,002,493 | 12,040,189 | 8,806,958 | 12,022,133 | 11,984,270 | 1,624,740 | 11,993,543 | 10,808,728 |
| Contig length N50 (bp) | 6,718,904 | 6,635,075 | 6,653,560 | 3,584,337 | 6,640,858 | 6,624,865 | 425,111 | 4,648,208 | 6,376,888 |
| L50 | 4 | 4 | 4 | 6 | 4 | 4 | 41 | 5 | 4 |
| Contig length N75 (bp) | 3,849,551 | 3,390,378 | 3,817,659 | 2,159,680 | 2,980,984 | 3,316,182 | 264,690 | 3,250,472 | 2,991,151 |
| L75 | 7 | 8 | 7 | 12 | 8 | 8 | 78 | 9 | 9 |
| GC (%) | 52.53 | 52.38 | 52.31 | 52.44 | 52.37 | 52.39 | 52.27 | 52.29 | 54.66 |

**Table S5. Comparison of the chromosome size between long-read assemblies and reference genomes.**

| **ToxoDB-48_GT1** | | | ***Tg*RH88 assembly** | | |  |
| --- | --- | --- | --- | --- | --- | --- |
| **Chrom-osome** | **Chr length (bp)** | **Telo-meres** | **Contig** | **Contig length (bp)** | **Difference (bp)** | **Telo-meres** |
| ChrIa | 1,841,710 | 1/2 | TgRH88_tig00000106_chrIa | 1,905,105 | 63,395 | 1/2 |
| ChrIb | 1,949,725 | 0/2 | TgRH88_tig00000006_chrIb | 2,036,895 | 87,170 | 2/2 |
| ChrII | 2,257,027 | 0/2 | TgRH88_tig00000031_chrII | 2,517,460 | 260,433 | 1/2 |
| ChrIII | 2,361,114 | 0/2 | TgRH88_tig00000013_chrIII | 2,653,274 | 292,160 | 1/2 |
| ChrIV | 2,527,423 | 0/2 | TgRH88_tig00000014_chrIV | 2,784,263 | 256,840 | 2/2 |
| ChrV | 3,030,196 | 0/2 | TgRH88_tig00000029_chrV | 3,482,671 | 452,475 | 2/2 |
| ChrVI | 3,539,197 | 0/2 | TgRH88_tig00000022_chrVI | 3,884,487 | 345,290 | 2/2 |
| ChrVIIa | 4,380,543 | 0/2 | TgRH88_tig00000012_chrVIIa | 4,643,533 | 262,990 | 2/2 |
| ChrVIIb | 4,975,702 | 0/2 | N/A | N/A | N/A | 2/2 |
| ChrVIII | 6,899,611 | 0/2 | TgRH88_tig00000030_chrVIII | 12,055,564 | 180,251 | 2/2 |
| ChrIX | 6,105,434 | 0/2 | TgRH88_tig00000001_chrIX | 6,606,532 | 501,098 | 1/2 |
| ChrX | 7,353,086 | 0/2 | TgRH88_tig00000003_chrX | 7,796,787 | 443,701 | 1/2 |
| ChrXI | 6,541,237 | 1/2 | TgRH88_scf00000009_chrXI | 6,778,623 | 237,386 | 2/2 |
| ChrXII | 6,851,637 | 0/2 | TgRH88_tig00000025_chrXII | 7,343,238 | 491,601 | 1/2 |
| Total | 60,613,642 | 12, 2, 0 | Total | 64,488,432 | 3,874,790 | 0, 6, 8 |
| **ToxoDB-48_ME49** | | | ***Tg*ME49 assembly** | | |  |
| **Chrom-osome** | **Chr length (bp)** | **Telo-meres** | **Contig** | **Contig length (bp)** | **Difference (bp)** | **Telo-meres** |
| ChrIa | 1,859,933 | 1/2 | TgME49_tig00000028_chrIa | 1,997,081 | 137,148 | 2/2 |
| ChrIb | 1,955,354 | 2/2 | TgME49_tig00000032_chrIb | 1,982,890 | 27,536 | 2/2 |
| ChrII | 2,347,032 | 0/2 | TgME49_tig00000022_chrII | 2,497,824 | 150,792 | 2/2 |
| ChrIII | 2,532,871 | 2/2 | TgME49_tig00000025_chrIII | 2,591,259 | 58,388 | 2/2 |
| ChrIV | 2,686,605 | 0/2 | TgME49_tig00000020_chrIV | 2,830,561 | 143,956 | 2/2 |
| ChrV | 3,331,915 | 0/2 | TgME49_tig00000016_chrV | 3,596,493 | 264,578 | 2/2 |
| ChrVI | 3,656,745 | 1/2 | TgME49_tig00012832_chrVI | 3,781,463 | 124,718 | 2/2 |
| ChrVIIa | 4,541,629 | 2/2 | TgME49_tig00000014_chrVIIa | 3,410,322 | 264,392 | 2/2 |
|  |  |  | TgME49_tig00000010_chrVIIa | 1,395,699 |  |  |
| ChrVIIb | 5,069,724 | 1/2 | N/A | N/A | N/A | 2/2 |
| ChrVIII | 6,970,285 | 1/2 | TgME49_tig00000001_chrVIII | 12,088,238 | 48,229 | 2/2 |
| ChrIX | 6,327,655 | 1/2 | TgME49_tig00000011_chrIX | 6,603,527 | 275,872 | 2/2 |
| ChrX | 7,486,190 | 0/2 | TgME49_tig00012828_chrX | 7,667,134 | 180,944 | 2/2 |
| ChrXI | 6,623,461 | 1/2 | TgME49_tig00012830_chrXI | 6,675,137 | 51,676 | 2/2 |
| ChrXII | 7,094,428 | 1/2 | TgME49_tig00000006_chrXII | 7,423,918 | 329,490 | 2/2 |
| Total | 60,623,894 | 4,7,3 | Total | 62,544,465 | 1,920,571 | 0,0,14 |
| **ToxoDB-48_VEG** | | | ***Tg*CTG assembly** | | |  |
| **Chrom-osome** | **Chr length (bp)** | **Telo-meres** | **Contig** | **Contig length (bp)** | **Difference (bp)** | **Telo-meres** |
| ChrIa | 1,874,844 | 1/2 | TgCTG_tig00000034_chrIa | 1,900,375 | 25,531 | 1/2 |
| ChrIb | 1,974,527 | 1/2 | TgCTG_tig00000018_chrIb | 1,977,238 | 2,711 | 2/2 |
| ChrII | 2,341,721 | 1/2 | TgCTG_tig00000001_chrII | 2,516,400 | 174,679 | 2/2 |
| ChrIII | 2,499,703 | 0/2 | TgCTG_tig00000014_chrIII | 2,588,969 | 89,266 | 2/2 |
| ChrIV | 2,697,628 | 1/2 | TgCTG_tig00000019_chrIV | 2,847,039 | 149,411 | 2/2 |
| ChrV | 3,263,790 | 0/2 | TgCTG_tig00000015_chrV | 3,431,465 | 167,675 | 2/2 |
| ChrVI | 3,680,711 | 0/2 | TgCTG_tig00000010_chrVI | 3,832,413 | 151,702 | 2/2 |
| ChrVIIa | 4,652,733 | 0/2 | TgCTG_tig00000011_chrVIIa | 4,684,986 | 32,253 | 2/2 |
| ChrVIIb | 5,093,952 | 0/2 | N/A | N/A | N/A | 2/2 |
| ChrVIII | 6,955,806 | 0/2 | TgCTG_tig00000012_chrVIII | 12,092,387 | 42,629 | 2/2 |
| ChrIX | 6,372,921 | 0/2 | TgCTG_tig00000007_chrIX | 6,607,067 | 234,146 | 2/2 |
| ChrX | 7,456,791 | 0/2 | TgCTG_tig00000004_chrX | 7,760,128 | 303,337 | 2/2 |
| ChrXI | 6,626,631 | 1/2 | TgCTG_tig00000020_chrXI | 6,680,781 | 54,150 | 2/2 |
| ChrXII | 7,144,635 | 0/2 | TgCTG_tig00000013_chrXII | 7,389,637 | 245,002 | 2/2 |
| Total | 62,636,393 | 9, 5, 0 | Total | 64,308,885 | 1,672,492 | 0, 1, 13 |

**Table S6: Data file containing updated copy number and coordinates for tandem gene arrays identified in *T. gondii* and *N. caninum***

**Table S7. Summary of the duplicated loci resolved by long-read assembly (loci are described in Adomako-Ankomah *et al.* (2014))**

| **Locus** | **Geno-type** | **Gene ID** | **Chr** | **Assembly** | **Copy number** | **Copy order** |
| --- | --- | --- | --- | --- | --- | --- |
| ROP5 | 1 | GenBank: HQ916451, HQ916448, HQ916455 | XII | *Tg*RH88 | 7 | c-c-c-b-a-b-a |
|  | 2 | GenBank: HQ916453, HQ916449, HQ916457 | XII | *Tg*ME49 | 9 | ND |
|  | 3 | GenBank: HQ916454, HQ916450, HQ916459 | XII | *Tg*CTG | 4 | c-c-b-a |
|  | 2 | GenBank: HQ916453, HQ916449, HQ916457 | XII | *Tg*CL13 | 9 | ND |
|  | 2 | GenBank: HQ916453, HQ916449, HQ916457 | XII | *Tg*S27 | 6 | ND |
|  | 2 | GenBank: HQ916453, HQ916449, HQ916457 | XII | *Tg*S21 | 6 | ND |
|  | 2 | GenBank: HQ916453, HQ916449, HQ916457 | XII | *Tg*S26 | 7 | ND |
|  | 3 | GenBank: HQ916454, HQ916450, HQ916459 | XII | *Tg*D3X1 | 4 | c-c-b-a |
|  | - | ToxoDB: NCLIV_060730 | - | *Nc*Liv | 2 | a-b |
| ROP38 | 1 | ToxoDB: TGGT1_242100 | VI | *Tg*RH89 | 2 | ND |
|  | 2 | ToxoDB: TGME49_242110 | VI | *Tg*ME50 | 3 | ND |
|  | 3 | GenBank: CEL73474 | VI | *Tg*CTG | 6 | ND |
|  | 2 | ToxoDB: TGME49_242110 | VI | *Tg*CL14 | 3 | ND |
|  | 2 | ToxoDB: TGME49_242110 | VI | *Tg*S27 | 3 | ND |
|  | 2 | ToxoDB: TGME49_242110 | VI | *Tg*S21 | 3 | ND |
|  | 3 | GenBank: CEL73474 | VI | *Tg*S26 | 2 | ND |
|  | 2 | ToxoDB: TGME49_242110 | VI | *Tg*D3X2 | 3 | ND |
|  | - | ToxoDB: NCLIV_017420 | - | *Nc*Liv | 4 | a-b-a-b |
| MIC17 | 1 | ToxoDB: TGGT1_200250, TGGT1_200240, TGGT1_200230 | VIII | *Tg*RH88 | 5 | a-b-c-c-c |
|  | 2 | ToxoDB: TGME49_200250, TGME49_200240, TGME49_200230 | VIII | *Tg*ME49 | 7 | a-b-c-c-c-c-c |
|  | 3 | ToxoDB: TGVEG_200250, TGVEG_200240 | VIII | *Tg*CTG | 7 | a-b-c-c-c-c-c |
|  | 3 | ToxoDB: TGVEG_200250, TGVEG_200240 | VIII | *Tg*CL13 | 7 | a-b-c-c-c-c-c |
|  | 3 | ToxoDB: TGVEG_200250, TGVEG_200240 | VIII | *Tg*S27 | 7 | a-b-c-c-c-c-c |
|  | 2 | ToxoDB: TGME49_200250, TGME49_200240, TGME49_200230 | VIII | *Tg*S21 | 7 | a-b-c-c-c-c-c |
|  | 2 | ToxoDB: TGME49_200250, TGME49_200240, TGME49_200230 | VIII | *Tg*S26 | 6 | a-b-c-c-c-c |
|  | 2 | ToxoDB: TGME49_200250, TGME49_200240, TGME49_200230 | VIII | *Tg*D3X1 | 7 | a-b-c-c-c-c-c |
|  | - | ToxoDB: NCLIV_038120, NCLIV_038110, NCLIV_038100, NCLIV_068830 | - | *Nc*Liv | 4 | a-d-b-c |
| MAF1 | 1 | GenBank: ANN02899, AMN92246, AMN92247, AMN92248, AMN92249 | II | *Tg*RH88 | 7 | b-a-b-a-b-a-c |
|  | 2 | GenBank: AMN92252, AMN92253 | II | *Tg*ME49 | 7 | b-a-b-a-b-a-c |
|  | 3 | GenBank: AMN92250, AMN92251 | II | *Tg*CTG | 5 | b-a-b-a-c |
|  | 2 | GenBank: AMN92252, AMN92253 | II | *Tg*CL13 | 7 | b-a-b-a-b-a-c |
|  | 2 | GenBank: AMN92252, AMN92253 | II | *Tg*S27 | 7 | b-a-b-a-b-a-c |
|  | 3 | GenBank: AMN92250, AMN92251 | II | *Tg*S21 | 5 | b-a-b-a-c |
|  | 2 | GenBank: AMN92252, AMN92253 | II | *Tg*S261 | 7 | b-a-b-a-b-a-c |
|  | 2 | GenBank: AMN92252, AMN92254 | II | *Tg*D3X1 | 7 | b-a-b-a-b-a-c |
|  | - | ToxoDB: NCLIV_004730 | - | *Nc*Liv | 2 | b-b |
| ROP4/7 | 1 | ToxoDB: TGGT1_295125, TGGT1_295110 | Ia | *Tg*RH88 | 3 | ROP4-ROP7-ROP7 |
|  | 2 | GenBank: EU047558; ToxoDB: TGME49_295110 | Ia | *Tg*ME49 | 5 | ROP4-ROP7-ROP4-ROP7-ROP7 |
|  | 3 | ToxoDB: TGVEG_295125, TGVEG_295110 | Ia | *Tg*CTG | 6 | ROP4-ROP7-ROP4-ROP4-ROP7-ROP7 |
|  | 3 | ToxoDB: TGVEG_295125, TGVEG_295110 | Ia | *Tg*CL13 | 6 | ROP4-ROP7-ROP4-ROP4-ROP7-ROP7 |
|  | 3 | ToxoDB: TGVEG_295125, TGVEG_295110 | Ia | *Tg*S27 | 6 | ROP4-ROP7-ROP4-ROP4-ROP7-ROP7 |
|  | 2 | GenBank: EU047558; ToxoDB: TGME49_295110 | Ia | *Tg*S21 | 5 | ROP4-ROP7-ROP4-ROP7-ROP7 |
|  | 2 | GenBank: EU047558; ToxoDB: TGME49_295110 | Ia | *Tg*S26 | 5 | ROP4-ROP7-ROP4-ROP7-ROP7 |
|  | 3 | ToxoDB: TGVEG_295125, TGVEG_295110 | Ia | *Tg*D3X1 | 6 | ROP4-ROP7-ROP4-ROP4-ROP7-ROP7 |
|  | - | ToxoDB: NCLIV_001950, NCLIV_001970 | - | *Nc*Liv | 5 | ROP4-ROP4-ROP4-ROP4-ROP7 |
| TSEL8 | 1 | - | III | *Tg*RH88 | 2 | a-b |
|  | 2 | - | III | *Tg*ME49 | 3 | c-a-b |
|  | 3 | - | III | *Tg*CTG | 4 | a-a-b-c |
|  | 2 | - | III | *Tg*CL13 | 3 | c-a-b |
|  | 2 | - | III | *Tg*S27 | 3 | c-a-b |
|  | 3 | - | III | *Tg*S21 | 4 | a-a-b-c |
|  | 3 | - | III | *Tg*S26 | 4 | a-a-b-c |
|  | 2 | - | III | *Tg*D3X1 | 3 | c-a-b |
| ^1^*MAF1* locus in S26 assembly is at the edge of contig_204. | | | | | |  |

**Table S8: Data file containing tandem repeats with period sizes > 500 bp in the T. gondii ME49 Nanopore assembly, along with their copy number and position in T. gondii RH88 and T. gondii VEG.**

Adomako-Ankomah Y, Wier GM, Borges AL, Wand HE, Boyle JP. 2014. Differential locus expansion distinguishes *Toxoplasma*tinae species and closely related strains of *Toxoplasma gondii*. *mBio* **5**: e01003-01013.

Brooks CF, Francia ME, Gissot M, Croken MM, Kim K, Striepen B. 2011. *Toxoplasma gondii* sequesters centromeres to a specific nuclear region throughout the cell cycle. *Proceedings of the National Academy of Sciences of the United States of America* **108**: 3767-3772.
